# Multiplexed fractionated proteomics reveals synaptic factors associated with cognitive resilience in Alzheimer’s Disease

**DOI:** 10.1101/2020.07.31.230680

**Authors:** B.C. Carlyle, S.E. Kandigian, J. Kreuzer, S. Das, B.A. Trombetta, Y. Kuo, D.A. Bennett, J.A. Schneider, V.A. Petyuk, R.R Kitchen, R. Morris, A.C. Nairn, B.T. Hyman, W. Haas, S.E. Arnold

## Abstract

Alzheimer’s disease (AD) is a complex neurodegenerative disease defined by the presence of amyloid-β (Aβ) plaques and tau neurofibrillary tangles, and driven by dysproteostatis, inflammation, metabolic dysfunction, and oxidative injury, eventually leading to synapse loss and cell death. Synapse loss correlates with cognitive impairment and may occur independently of the extent of AD pathology. To understand how synaptic composition is changed in relation to AD neuropathology and cognition, highly sensitive multiplexed liquid chromatography mass-spectrometry was used to quantify biochemically enriched synaptic proteins from the parietal association cortex of 100 subjects with contrasting AD pathology and cognitive performance. Functional analysis showed preservation of synaptic signaling, ion transport, and mitochondrial proteins in normal aged and “resilient” (cognitively unimpaired with AD pathology) individuals. Compared to these individuals, those with cognitive impairment showed significant metabolic differences and increased immune- and inflammatory-related proteins, establishing the synapse as a potential integration point for multiple AD pathophysiologies.

## Introduction

Alzheimer’s disease (AD) is the most common cause of dementia, affecting an estimated 5.7 million Americans in 2018 and about 35 million individuals worldwide (Alzheimers & Dementia, 2018). Abundant amyloid-β (Aβ) plaques and paired helical filament tau (PHF-tau) neurofibrillary tangles in the cerebral cortex define the disease neuropathologically, but there is increasing recognition that the level of cognitive impairment associated with these pathologies is variable. Some individuals exhibit no discernable impairments during their lifetime, despite having high levels of AD pathology. This phenomenon has been described in both autopsy and biomarker studies of aging and AD and is variously referred to as resilience, reserve, asymptomatic AD, or preclinical AD (Arnold et al., 2013; Au et al., 2012; D. A. Bennett et al., 2006; Gelber, Launer, & White, 2012; Iacono et al., 2009; O’Brien et al., 2009; Savva et al., 2009; Schneider, Arvanitakis, Bang, & Bennett, 2007). On the other hand, some individuals show more severe impairments than might be expected in the setting of minimal AD pathology, cerebrovascular disease, or other neurodegenerative diseases in the brain. This cognitive frailty is less well-studied. The relationship between AD pathology and cognitive impairment may weaken yet further in individuals over 90 years of age (Ewbank & Arnold, 2009; Haroutunian et al., 2008). Understanding the cellular and molecular basis of brain response to AD and other common pathologies of aging is of paramount importance for prevention and treatment discovery. Synapse loss has long been considered as a strong correlate of cognitive impairment in AD (DeKosky & Scheff, 1990; Koffie, Hyman, & Spires-Jones, 2011; Terry et al., 1991). Clinicopathological studies of individuals enrolled in the Religious Orders Study and Memory and Aging Project (ROSMAP) (David A Bennett et al., 2018) cohorts highlighted comparable pre-synaptic and post-synaptic staining levels in normal controls and resilient individuals. Another study using dendrite tracing in the dorsolateral prefrontal cortex suggested resilient individuals have a similar density of thin and mushroom spines relative to controls (Boros et al., 2017). A recent proteomic study of post-mortem tissue from the frontal cortex and anterior cingulate of individuals with AD showed a decrease in the pre-synaptic markers SNAP25, Syntaxin 1A & B (STX1A & B), and synaptotagmin (SYT1), and the post-synaptic markers PSD95, disks large MAGUK scaffold protein 3 (DLG3), and Neuroligin 2 (NLGN2) (Ping et al., 2018). A further recent study of cognitive trajectory over time showed that aged individuals with a worse cognitive trajectory had lower levels of synaptic markers, including PSD95, SYT1, and STX1A in post-mortem tissues (Wingo et al., 2019).

The challenge of interpreting whole-tissue data with regards to protein differences in specific cellular compartments in a cytoarchitecturally complex tissue like the brain is that measurements may be more sensitive to global differences in organelle density or volume than organelle composition itself (Becky C. Carlyle et al., 2017). In whole tissue studies of AD, one of the strongest drivers of differential protein abundance is the general loss of synaptic markers in Dementia-AD cases compared to controls (Johnson et al., 2018a; Ping et al., 2018; Wingo et al., 2019). Therefore, to better understand how protein composition at the synapse is affected in relation to AD pathology and cognition, we analyzed enriched synaptic fractions in brain tissue from ROSMAP by proteomic tandem mass tag labelled liquid chromatography mass-spectrometry (LC-MS3). Parietal association cortex (angular gyrus) tissue from 100 participants spanning four groups was analyzed: 1) cognitively unimpaired with low AD or other pathology (“Normal”), 2) dementia with abundant AD pathology (“Dementia-AD”), 3) “Resilient,” defined as cognitively unimpaired despite abundant AD pathology, and 4) “Frail,” defined as dementia without AD or any other attributable pathology.

We identified functional clusters of proteins that were enriched in cognitively impaired individuals versus non-impaired and resilient individuals. These upregulated categories included metabolic, extracellular matrix (ECM) remodeling, and immune and inflammation functions. Previous studies have suggested that these differences may arise mostly from non-neuronal cell types (Johnson et al., 2018a) but in this work we show that the synapse may also act as a nexus between these processes and cognitive impairment. Cognitively resilient individuals showed more mitochondrial and synaptic signaling proteins in the absence of a tissue volume effect. These proteins may act as cognitively relevant biomarkers of synaptic function in AD and offer novel pathways for therapeutic targeting.

## Results

### Sample demographics of diagnostic groups

The 100 samples were systematically selected from the ROSMAP cohorts to represent contrasting degrees of disease pathology and cognitive impairment. All had detailed demographic, clinical, psychometric, and neuropathological data from the ROSMAP studies (David A Bennett et al., 2018)(TableS1). Samples were classified into four diagnostic groups on the basis of two variables; the Braak score (1-4, low AD pathology, 5-6, high AD pathology) and clinical consensus of the presence of significant cognitive impairment at last study visit prior to death (Figure 1A). Twenty-five samples were allocated to each of four groups; Normal individuals with low AD pathology and no cognitive impairment, Dementia-AD individuals with high AD pathology and cognitive impairment, Resilient individuals with high AD pathology and no cognitive impairment, and Frail individuals with low AD pathology and cognitive impairment. Key sample demographics were well matched across the 4 diagnostic categories including age at death, post-mortem interval (Figure 1B), sex and education (Table 1).

**Table 1:**
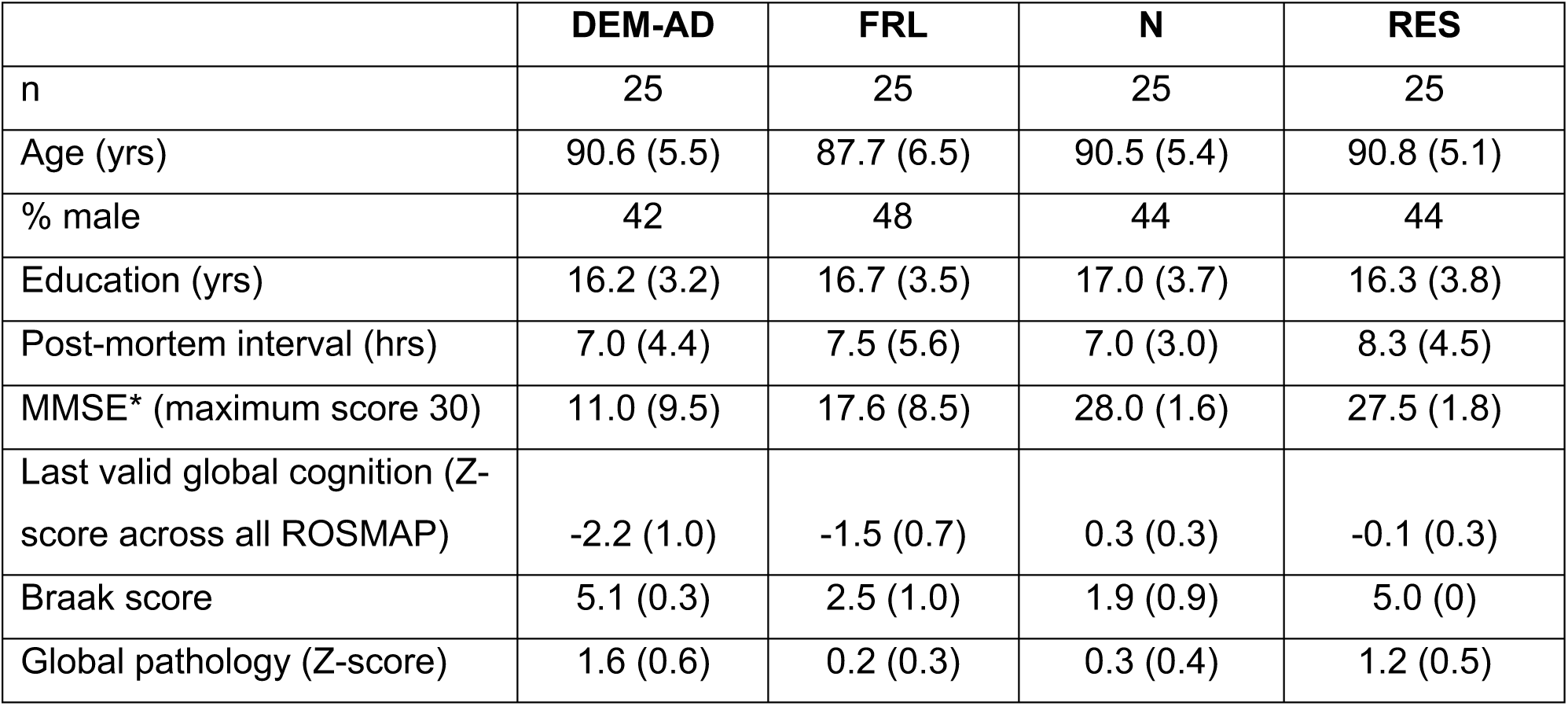
Summary demographics of cases. Data is presented as mean (standard deviation). See Supplementary Table 1 for individual sample demographic data. * MMSE = Mini Mental State Examination

**Figure 1:**
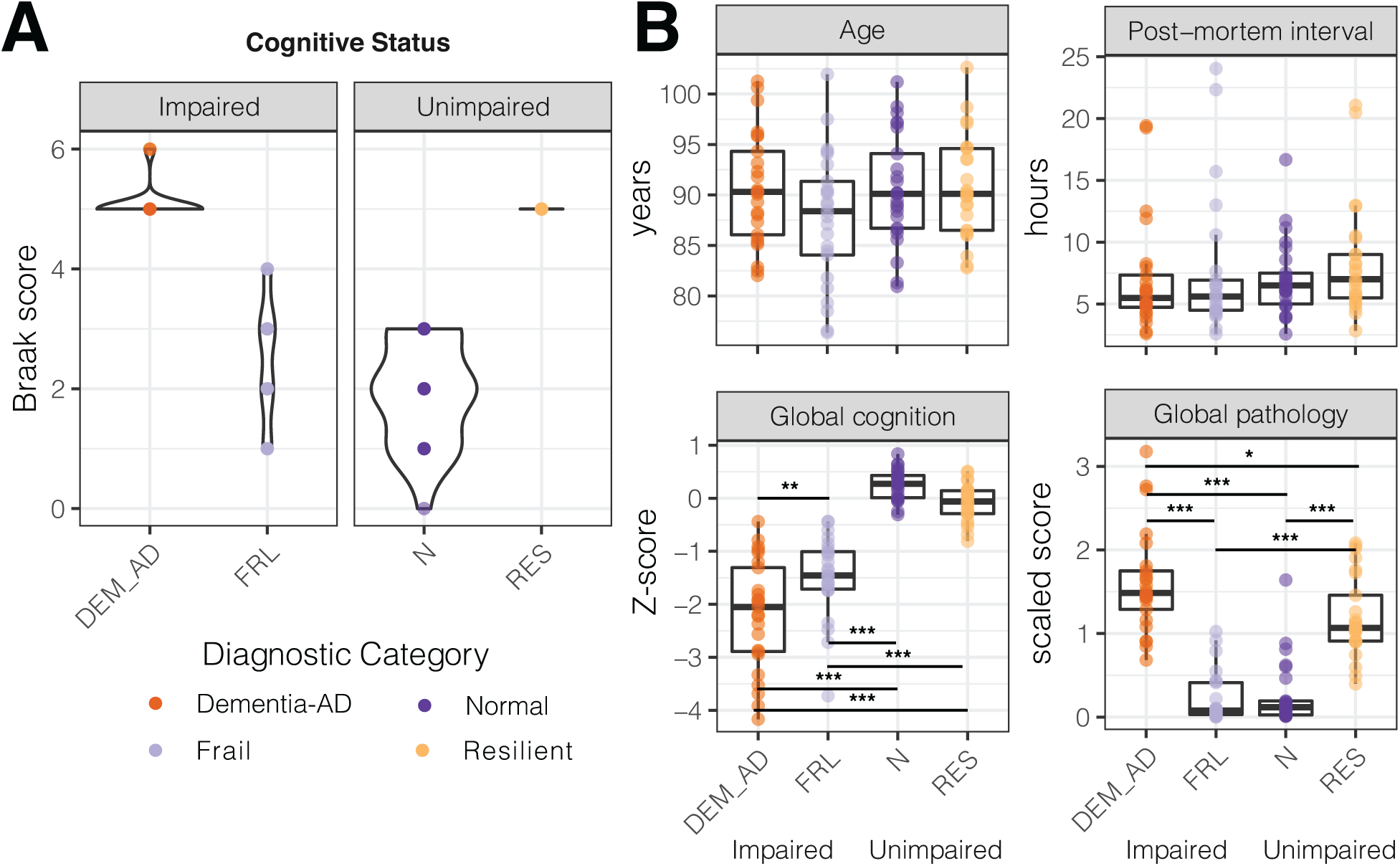
One hundred subject samples were categorized based on their levels of AD pathology and cognitive performance. **A)** Samples were divided into 4 groups by Braak score (Braak > 4 high pathology, Braak ≤ 4 low pathology) and clinical diagnosis of the presence of dementia-level cognitive impairment (n = 25 per group). **B)** Age and post-mortem interval were well balanced across all diagnostic groups. Global cognition score at last valid visit was significantly higher in Normal and Resilient subjects compared to Dementia-AD and Frail subjects. There was a small but significant difference in global cognition score between Dementia-AD and Frail subjects. Global pathology score (a scaled composite score accounting for diffuse and neuritic plaques and neurofibrillary tangles) was significantly higher in Dementia-AD and Resilient subjects than Normal and Frail Subjects. There was a small but significant difference in global pathology between Dementia-AD and Resilient subjects. Between group significance was defined by one-way ANOVA followed by Tukey post-hoc testing, (*adjusted p < 0.05, **p < 0.01,***p < 0.001)

One-way ANOVA with Tukey post-hoc testing showed that the global pathology score was significantly higher in the Dementia-AD and Resilient groups than the Normal and Frail groups (Figure 1B), and that the global cognition score at the last valid visit was significantly higher in the Normal and Resilient groups compared to the Frail and Dementia-AD groups (Figure 1B, Table S2). There was also a smaller significant difference in global cognition score between the Frail and Dementia-AD groups. These subjects were not selected for balanced ApoE status across groups, and thus this is not used as a variable in the downstream analysis. ApoE4 risk allele carrier distributed as expected across the groups; 8% of Normal subjects, 24% of Frail and Resilient subjects, and 44% of Dementia-AD subjects (Figure S1). There were zero carriers of the protective ApoE2 variant in the Dementia-AD group, 28% in the Normal group, 32% in the Frail group, and 8% in the Resilient group. Although this variable was not included in the modeling, single protein data can be explored relative to ApoE status at https://tmt-synaptosomes.omics.kitchen/. Lewy body pathology was absent in all but four Resilient subjects, and vascular macroinfarcts were also present only in 9 of the Resilient group. Vascular microinfarcts were more widely present throughout the groups, being present in 16% of Dementia-AD subjects, 32% of Frail and Resilient subjects, and 8% of Normal subjects. Similarly there were no differences in TDP-43 pathology across the four groups, although there were 16 missing data points for this variable (Figure S2).

### Quantitative assessment of synaptic proteomes

Frozen tissue sections were obtained from the parietal association cortex (Brodmann area 39, angular gyrus). Synaptic proteins were enriched from approximately 100 mg of each tissue sample using the Syn-PER Synaptic Protein Extraction Reagent, which uses non-denaturing cell lysis to release organelles. P2 pellets were Tandem Mass Tag (TMT) labeled and prepared for analysis by LC-MS3 in 10 batches of 11 samples (Figure S3). In each batch the 11^th^ sample was a pooled common sample used for batch-to-batch normalization. Prior to LC-MS3 analysis, each 11-plex was offline fractionated into 12 fractions by basic Reverse Phase Liquid Chromatography (bRPLC) to ensure deep coverage of the synaptic proteome. Quantitative profiles for each of the 100 synaptic protein samples were acquired using multiplexed proteomics by applying TMT technology on an Orbitrap Fusion mass spectrometer using the SPS-MS3 method (McAlister et al., 2012, 2014; Ting, Rad, Gygi, & Haas, 2011). MS2 level peptide spectra were assigned to peptides and proteins using the Sequest algorithm (Eng, McCormack, & Yates, 1994), with two step normalization, protein level quantification, and upstream filtering performed using an in-house software suite (Huttlin et al., 2010a). Downstream analyses were performed in R, and the Gene Set Enrichment Analysis (Subramanian et al., 2005) software (Figure 2A).

**Figure 2:**
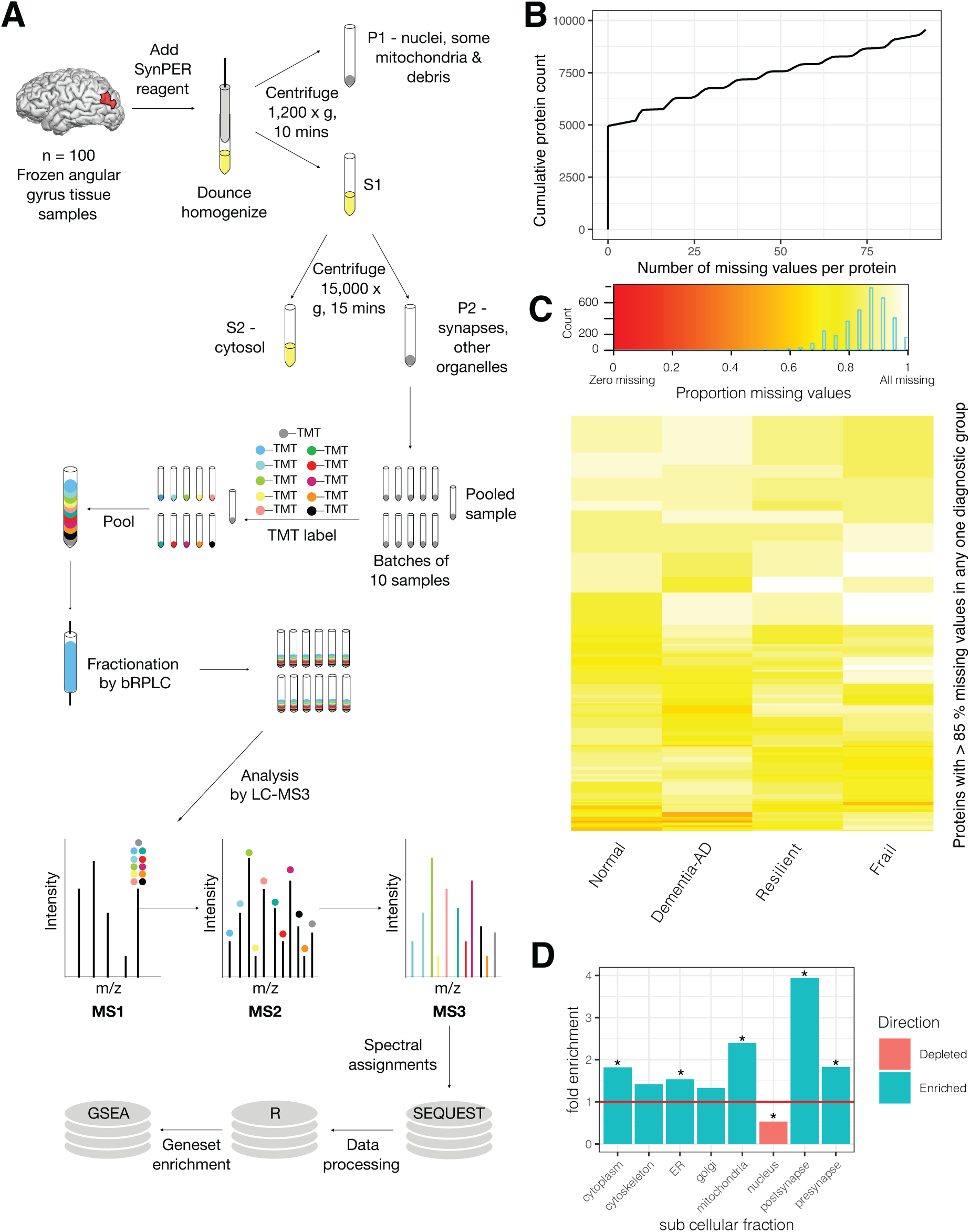
Fractionated LC-MS3 was used to assess synaptic protein enriched fractions from the angular gyrus. **A)** Schematic diagram showing experimental workflow. Cortical grey matter samples from the angular gyrus (BA 39) were fractionated to enrich for synaptic proteins using the SynPER reagent and low speed centrifugation. Synaptic protein fractions were TMT labelled, pooled, and offline fractionated into twelve fractions. Fractions were analyzed by LC-MS3. Spectra were assigned using SEQUEST and quantified using a custom software pipeline. Data analysis and figures were prepared in R and using the GSEA java applet. **B)** 9560 proteins were detected in at least one sample across the experiment. 4952 of these proteins were detected in every sample (ie. no missing values on the x-axis), with the stepped structure of this plot suggesting that protein detection generally followed the experimental batching structure. **C)** There were no proteins with a majority of missing values in one diagnostic group that were consistently detected (>55%) in one of the other diagnostic groups. The 872 proteins with over 85% of values missing in any one diagnostic group are visualized in a heatmap according to the proportion of missing values in each diagnostic group. **D)** Comparison of detected proteins with consensus lists of unique cellular fraction-associated protein IDs shows strong enrichment of the appropriate cellular compartments, with substantial depletion of the nuclear fraction. Maximum possible fold enrichment in this experiment was 4.2 fold (4874 unique GeneIDs observed from a possible 20,635), with the post-synaptic fraction having a fold enrichment of 3.9. (*Fisher test, Bonferroni adjusted p value < 0.001).

Across all non-pooled samples, 9560 unique proteins were detected and quantified in at least one subject sample. 4952 proteins were detected in every sample and for the remaining proteins, detection was mostly related to batching structure (Figure 2B). No proteins which were detected consistently (> 45% of the time) in one or more diagnostic groups but not in other diagnostic groups (Figure 2C). Due to the restricted nature of the cellular compartment being evaluated in this experiment, and to avoid difficulties with imputing missing values, the batch and TMT-label median normalized dataset was filtered to retain only those proteins quantified in every sample. Plotting of individual samples showed successful batch by batch median normalization (Figure S4A) but relatively variable distributions at quantification extremes. Values were therefore quantile normalized and samples clustered for visual inspection. No clear batch effects were evident across the samples from this clustering (Figure S4B).

### Enrichment and coverage of the synaptic proteome

The Syn-PER kit was chosen for synaptic fraction enrichment for two reasons; 1) Once tissue is frozen without cryopreservation methods, membrane disruption prevents the preparation of pure tissue fractions (Dias, Gandra, Brenzikofer, & Macedo, 2020) and 2) the protocol is simple and rapid compared to density gradient methods, reducing the potential introduction of variability from sample preparation methods in a large sample set. While the Syn-PER kit had been tested in-house for reproducible enrichment of the appropriate cell fraction by western blotting of a small number of marker proteins (data not shown), a more global view of appropriate cell fraction enrichment can be gained from the proteins detected across all samples in this experiment. Cell fraction protein consensus lists were prepared from proteins detected in a specific fraction in at least two published human or mouse tissue proteomic studies (Bayés et al., 2014, 2011; Christoforou et al., 2016; Distler et al., 2014; Föcking et al., 2016; Foster et al., 2006; Itzhak et al., 2017; Li et al., 2017; Pirooznia et al., 2012; Thul et al., 2017) or gene ontology cellular compartment protein lists (Ashburner et al., 2000;

The Gene Ontology Consortium, 2019). The list for each fraction was then restricted to proteins that were unique to one fraction. Bonferroni corrected Fisher tests were performed to assess enrichment or depletion of organelles. All fractions tested, except the nucleus, were enriched (Figure 2D), although golgi and cytoskeleton enrichments were not significant. The fractions with the strongest enrichment were post synaptic (3.9-fold), and mitochondrial (2.4-fold, Table S2). There are many fewer proteomics studies of the pre-synapse than post-, and thus incomplete annotation of unique proteins may be part of the reason why this fraction appears less enriched in this analysis. All fractions found to be significantly enriched in this preparation are organelles with established presence in the pre- or post-synapse. As expected, the nucleus was significantly depleted from this preparation (p.adj = 2.84e^-30^).

### Biochemical enrichment of synaptic proteins effectively controls for synapse loss between diagnostic groups

In whole tissue proteomic studies, one of the strongest drivers of changes in the data is a general loss of synaptic markers in Dementia-AD cases compared to controls (Johnson et al., 2018a; Ping et al., 2018; Wingo et al., 2019). This is likely a reflection of the fact that synapses will occupy a decreased volume of the grey matter once synapse loss occurs. We removed this potential confound by biochemically enriching the synaptic fraction, to focus on intrinsic protein changes within existing synapses. Established pre- and post-synaptic markers were plotted and assessed for protein abundance differences that may indicate gross synapse loss between groups. None of the established synaptic markers assessed were significantly different between groups by one-way ANOVA (Figure 3A, 3B, Table S4). This shows that biochemical enrichment of synaptic proteins was effective in avoiding the potential confound of synapse loss, particularly in the Dementia-AD category.

**Figure 3:**
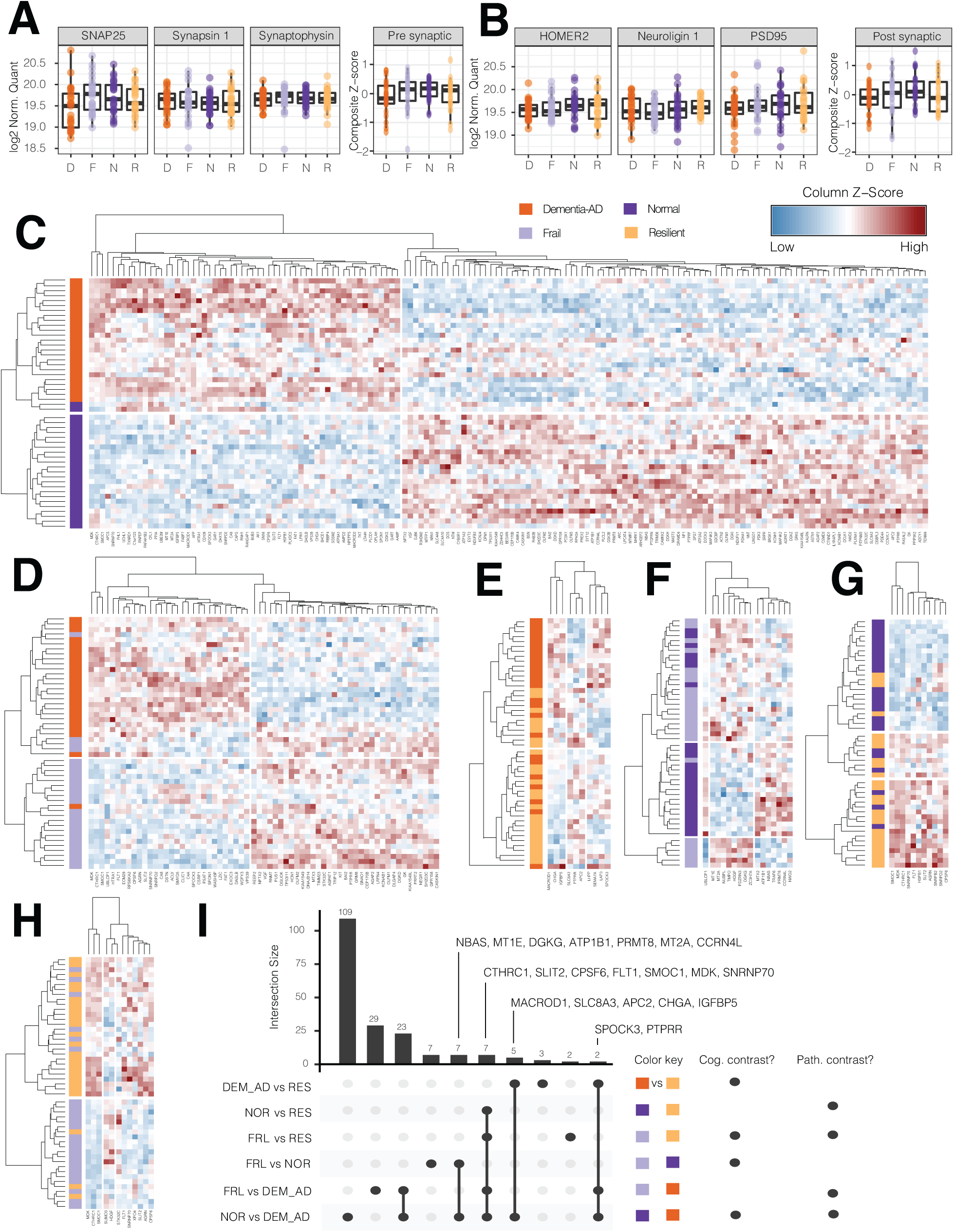
In total, 199 unique proteins were differentially expressed between diagnostic groups, with no clear sign of a volume artefact from synapse loss in Dementia-AD subjects. **A)** Abundance of established pre-synaptic markers across the four diagnostic groups suggested there was no significant volume artefact arising from gross synapse loss between groups. The pre-synaptic summary plot is a Z-score normalized composite of SYT1, NRZN3, SNAP25, SYN2, STX1A, STX1B, SLC17A6, SYN1, and SYP. **B)** Abundance of established post-synaptic markers across the diagnostic groups suggest there is no volume artifact arising from gross synapse loss between groups. The post-synaptic summary plot is a Z-score normalized composite of EPHB1, DLG3, DLG4 (PSD95), HOMER2, NLGN1, NLGN2, and SHANK1. **C)** Heatmap of differentially expressed proteins between Dementia-AD and Normal cases shows 156 proteins. Clustering on abundance of these 156 proteins produces almost perfect separation between Dementia-AD and Normal subjects. Heatmap of differentially expressed proteins between **D)** Dementia-AD and Frail, **E)** Dementia-AD and Resilient, **F)** Normal and Frail, **G)** Normal and Resilient, and **H)** Resilient and Frail subjects. **I)** Upset plot shows the intersection of proteins common to multiple comparisons. Seven proteins were common to all pathology contrasts, while no proteins were common to all cognitive contrasts.

### Categorical analysis of diagnosis and synaptic protein abundance

Linear models were constructed with the 4952 proteins present in all samples as outcome variables, and diagnostic category, age, gender, education and postmortem interval as explanatory variables. Summary tables were prepared using the R broom package, and p values were FDR corrected. FDR corrected p < 0.05 was considered significant. The group comparison with the largest number of significantly associated proteins is the highest contrast diagnostic group comparison, i.e., Dementia-AD versus Normal controls. 58 proteins are significantly increased in the Dementia-AD group versus Normal, and 98 are significantly decreased (summary Table 2, Table S5 for all protein data, Table S6 for significant only data, visualize individual protein plots at https://tmt-synaptosomes.omics.kitchen/). Unique to this contrast were established Alzheimer’s risk proteins APP and BACE1, and PDE4A, a phosphodiesterase enzyme previously linked to cognitive dysfunction in aging rodents and non-human primates (Becky C Carlyle et al., 2014). Clustering the Normal versus Dementia-AD samples on the basis of these proteins resulted in almost perfect separation of the two diagnostic groups (Figure 3C).

**Table 2:**
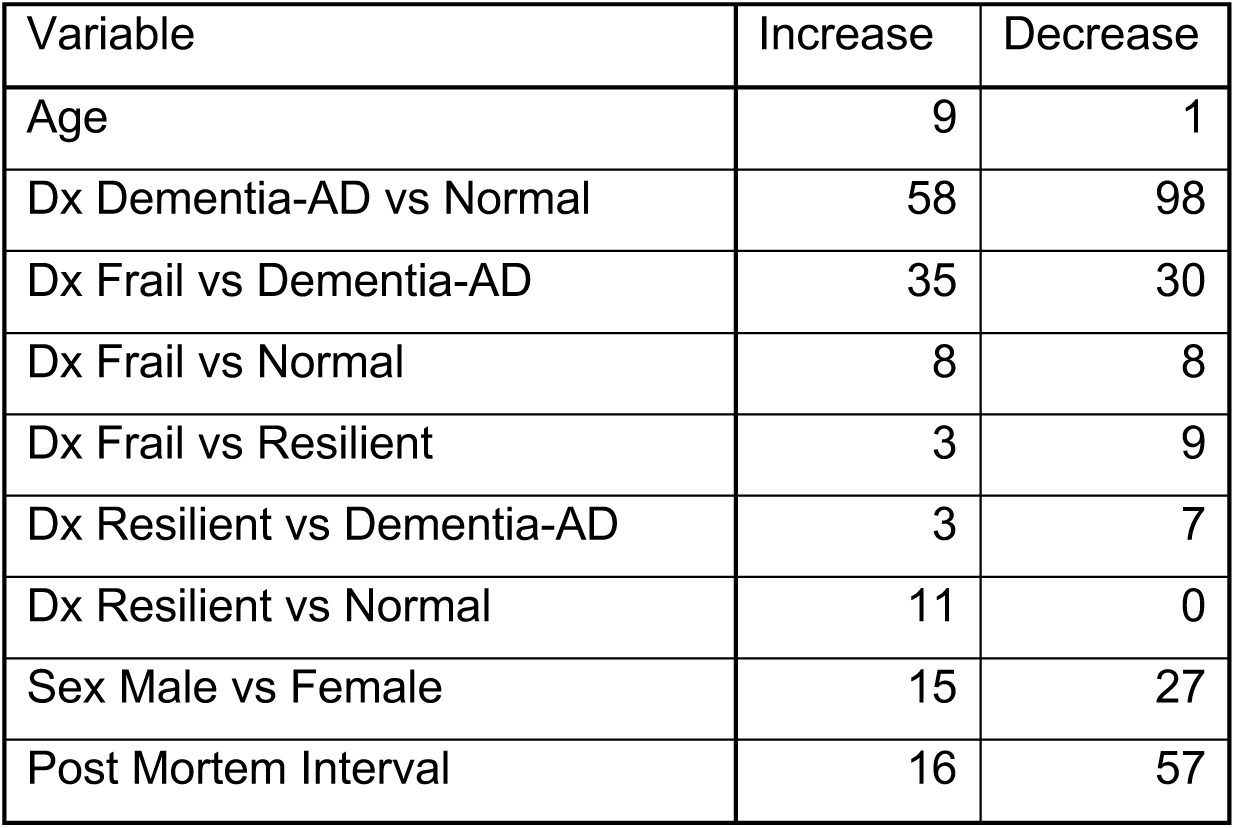
Summary of proteins significantly associated with each explanatory variable. No proteins were associated with level of education.

| Variable | Increase | Decrease |
| --- | --- | --- |
| Age | 9 | 1 |
| Dx Dementia-AD vs Normal | 58 | 98 |
| Dx Frail vs Dementia-AD | 35 | 30 |
| Dx Frail vs Normal | 8 | 8 |
| Dx Frail vs Resilient | 3 | 9 |
| Dx Resilient vs Dementia-AD | 3 | 7 |
| Dx Resilient vs Normal | 11 | 0 |
| Sex Male vs Female | 15 | 27 |
| Post Mortem Interval | 16 | 57 |

The comparison between the Dementia-AD and Frail groups was also striking, with clustering on the basis of these proteins providing good separation between the two groups. 35 proteins were upregulated in the Frail group and 30 were downregulated compared to Dementia-AD (Figure 3D). VGF and NPTX2, two of the most well replicated findings in proteomic studies of AD brain tissue, were found to be dysregulated in both the Frail and Normal versus Dementia contrasts. For the remaining comparisons there were fewer significant proteins (Table 2), and the clustering was less consistent between the two groups (Figure 3E, 3F, 3G, 3H).

Seven proteins were significantly associated with all comparisons where AD pathology is a contrast; CTHRC1, SLIT2, CPSF6, FLT1, SMOC1, MDK, and SNRNP70 (Figure 3I, 4A). All seven of these proteins were more abundant in the high pathology Dementia-AD and Resilient groups than the Normal and Frail groups. There were no proteins associated with all comparisons where cognitive status was a contrast. Seven proteins were shared between the Frail versus Normal and the Dementia-AD versus Normal comparisons; NBAS, MT1E, DGKG, ATP1B1, PRMT8, MT2A, and CCRN4L (Figure 3I, 4B). For most of these proteins levels were intermediate in Resilience, but variable enough that this was not significant. A further seven were shared between the Dementia-AD versus Resilient and Dementia-AD versus Normal comparisons; MACROD1, SLC8A3, APC2, CHGA, IGFBP5, SPOCK3, and PTPRR (Figure 3I, 4C). The lack of overlap in proteins significantly associated with cognition in individuals with or without AD pathology suggests that there may be two separate mechanisms involved; one in cognitive frailty in the absence of AD pathology and a second in cognitive resilience in the presence of AD pathology. This may explain why a two-way ANOVA (pathology*cognition) on these data does not show any proteins significantly associated with a main effect of cognition.

**Figure 4:**
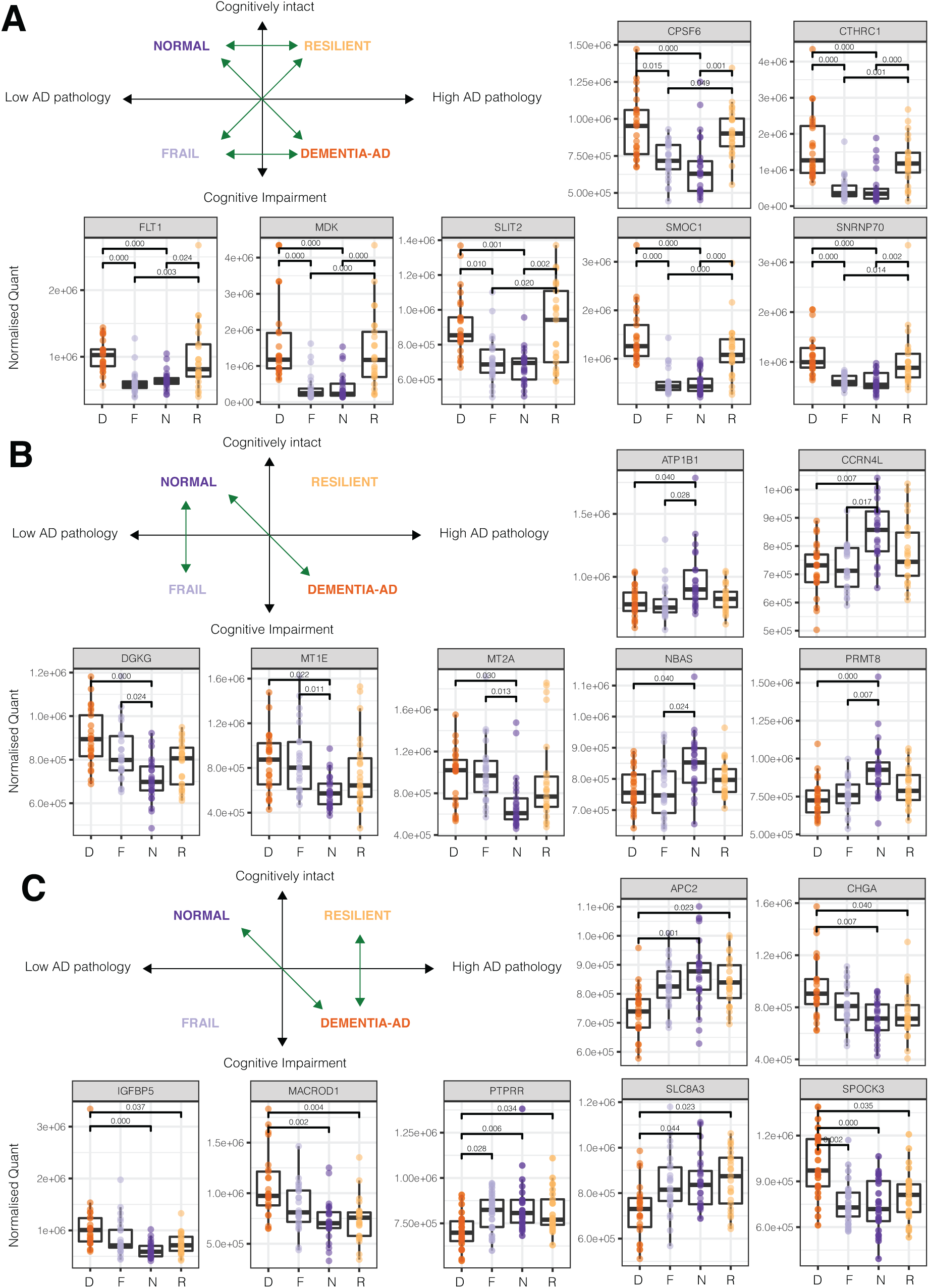
A small number of proteins tracked consistently with pathological contrasts or a subset of cognitive contrasts. **A)** Box plots to illustrate the seven proteins that were increased in every pathological contrast. Adjusted p-values are given to three significant figures. **B)** Box plots to illustrate proteins that were significant in both the Normal vs Dementia-AD and Normal versus Frail contrasts. **C)** Box plots to illustrate proteins that were significantly associated with both the Normal vs Dementia-AD and Dementia-AD versus Resilient contrast.

### Gene set enrichment analyses of Dementia-AD versus Normal samples

Non-parametric gene set enrichment analysis (GSEA) was used to examine functional differences between samples in the highest contrast sample set, the Dementia-AD versus Normal group. Using gene-set permutation, 284 Gene Ontology (GO) terms were significant (FDR q value less than 0.05, Table S7). For clarity of plotting, GO terms with a common parent were collapsed using ontological information from the GSA R package. Z-scores were calculated for each observed protein member of a GO term, and these Z-scores were averaged to produce a composite Z-score for each subject for each GO term. Clustering the Dementia-AD and Normal samples on the basis of these GO term Z-scores led to reasonably strong separation of samples by diagnostic category (Figure 5, non-collapsed version with all individual terms Figure S5). Terms representing metabolism, particularly NADH and NADP metabolism, innate and adaptive immune response, vascular endothelial growth factor signaling, and migration of immune cells were strongly enriched in Dementia-AD samples. Conversely, terms representing synaptic signaling, including Glutamate and GABA signaling, and mitochondrial oxidative phosphorylation were strongly enriched in the Normal samples (Figure 5).

**Figure 5:**
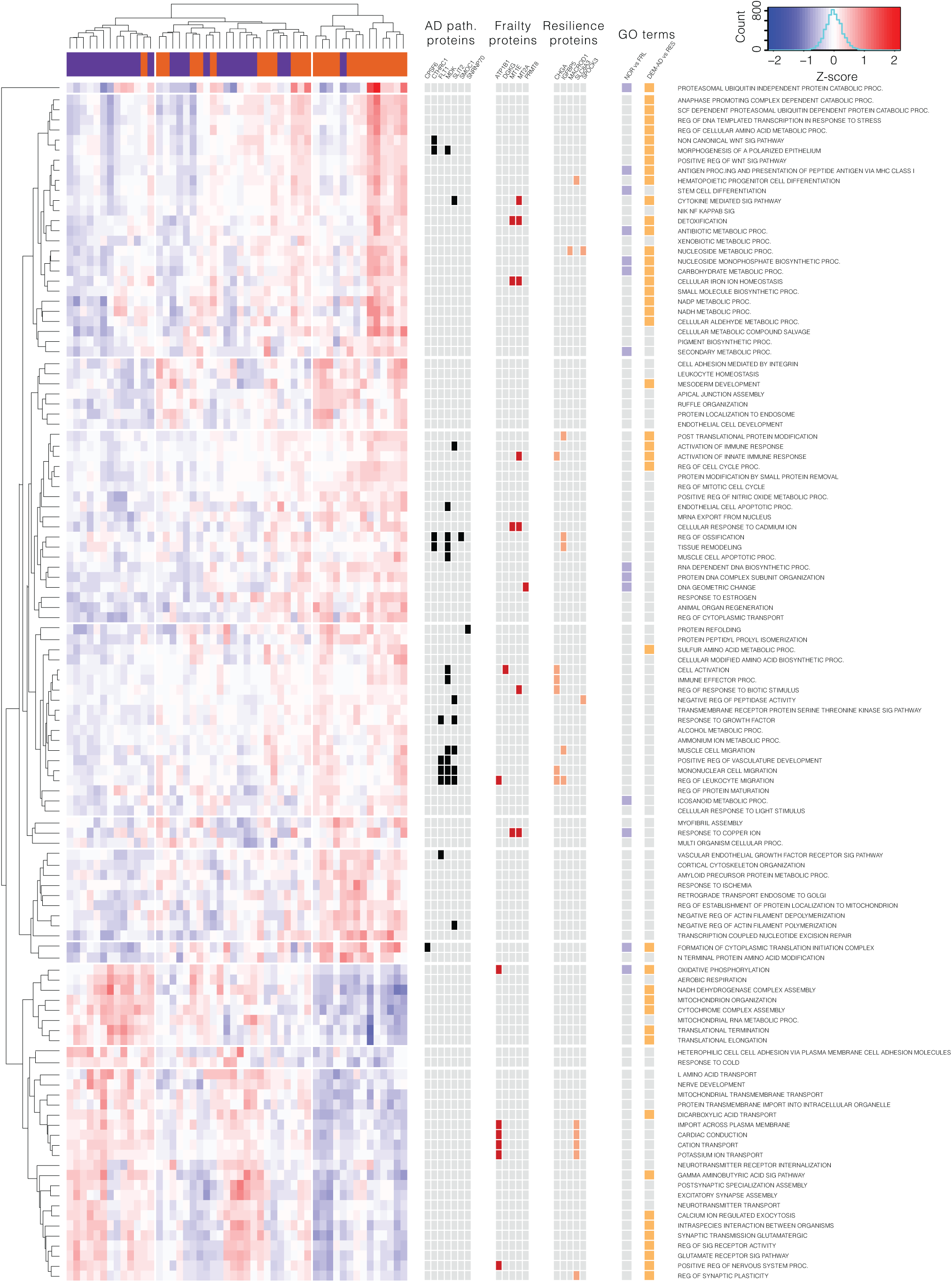
Gene Set Enrichment Analysis shows that synaptic proteins from Dementia-AD samples are enriched for GO terms involved in metabolism, extracellular matrix remodeling, and immune regulation. Synaptic proteins from Normal samples are enriched for synaptic signaling, mitochondrial and ion transport proteins. **A)** Heatmap shows FDR significant (p < 0.05) clustered GO-terms from the Dementia-AD versus Normal comparison, with related GO terms collapsed to the highest parent (See Figure SX for non-collapsed version). Significant proteins from Figure 4 are shown alongside, with the GO terms they belong to highlighted. The final boxes show which GO terms are shared between the other two cognitive contrasts. There is a greater overlap with Dementia-AD versus Resilience than with Normal vs Frail. Common words in GO terms have been shortened for plotting; Proc. = Process, Reg. = Regulation, Sig. = Signaling.

The 7 proteins involved in all AD pathology contrasts, with the exception of SLIT2, have not been studied extensively in the central nervous system. To learn more about their potential function, proteins were mapped to GO terms they were associated with. CTHRC1, MDK and SMOC1 clustered in the Regulation of Ossification and Tissue remodeling GO terms, suggesting a potential role in extra-cellular matrix (ECM) remodeling. FLT1, MDK, and SLIT2 are associated with a number of immune/inflammatory categories, particularly those related to migration of immune cells. The Frailty contrast (Significant in Normal vs Frail and Normal vs Dementia-AD) Na/K+ ATPase ATP1B1 was also associated with Regulation of Leukocyte Migration, in addition to established roles in ion transport. MT2A was associated with the terms Cytokine Mediated Signaling Pathway and Activation of Innate Immune Response. MT1E and MT2A are also associated with multiple terms involving response to metal ions. Resilience contrast (Significant in Normal vs Resilient and Normal vs Dementia-AD) protein CHGA is also associated with multiple immune terms, while IGFBP5 appears in multiple ECM related terms (Figure 5).

A table was produced to show which GO categories were populated by significantly differentially expressed proteins by direction of change (Table S8). In the Synaptic Signaling GO term, eighteen proteins were decreased in Dementia-AD compared to Normal subjects (ADCY1, ARC, ATP2A2, BRSK1, BSN, DGKI, DGKZ, GSK3B, IL1RAPL1, NF1, NPTX2, PLCL2, RAB3B, RPH3A, SDCBP, SLC4A10, SLC8A3, SYT12), with only three proteins enriched (known AD proteins APP and BACE1, plus DAGLB). Terms such as Cation Transport were also heavily biased in this direction, containing sixteen proteins that decreased in Dementia-AD (ACTN2, ARC, ATP1A3, ATP1B1, ATP2A2, IL1RAPL1, KCNAB1, KCNH1, NNT, PLCL2, SLC4A10, SLC8A3, SPG7), and four that increased (APP, ATP8A1, FHL1, PLCD1). In the opposite direction, Immune Effector Process contained eight proteins that were upregulated in Dementia-AD (APCS, APP, ATP8A1, CHGA, HTRA1, LTA4H, MDK, PPIA) versus two that were downregulated (PLCL2, SDCBP), and Response to Growth Factor contained seven upregulated proteins (APP, CD109, FLT1, HSPB1, HTRA1, SLIT2, SNX6) versus one down (SDCBP).

### Gene Set Enrichment analysis of Cognitively Contrasted Samples

To identify GO terms that were specifically related to cognitive impairment, in the presence or absence of pathology, gene set enrichment analysis was also performed on the two diagnostic comparisons that were matched for AD pathology, but divergent for cognitive performance: the Frail versus Normal groups and the Resilient versus Dementia-AD groups. Despite the low or absent levels of AD pathology in the Frail group, 43 GO terms were significant (FDR q value less than 0.05, Table S9) in the Frail versus Normal comparison. All significant GO terms were enriched in the Frail samples compared to the Normal samples (Figure 6B). The majority of GO terms (28 terms) were unique to this contrast, while fourteen overlapped with the Dementia-AD versus Normal contrast and five with the Dementia-AD vs Resilient contrast (Figure 6A). There are a small number of immune and inflammatory terms in this contrast, and a more substantial representation in this contrast of terms associated with DNA and RNA metabolism, chromatin organization, and splicing, which we hypothesized may reflect closer engagement of dividing cells such as microglia or astrocytes with the synaptic compartment. However this was not reflected in the dataset, where astrocytic markers GFAP, ALDH1L1, and GLUL were present in all samples in this dataset, as was the microglial marker CD11b (ITGAM), but there was no significant difference between diagnostic categories in these markers (Table S5). There is not a substantial overlap of significant differentially abundant proteins with informative GO terms in this contrast (Figure 6B, Table S10).

**Figure 6:**
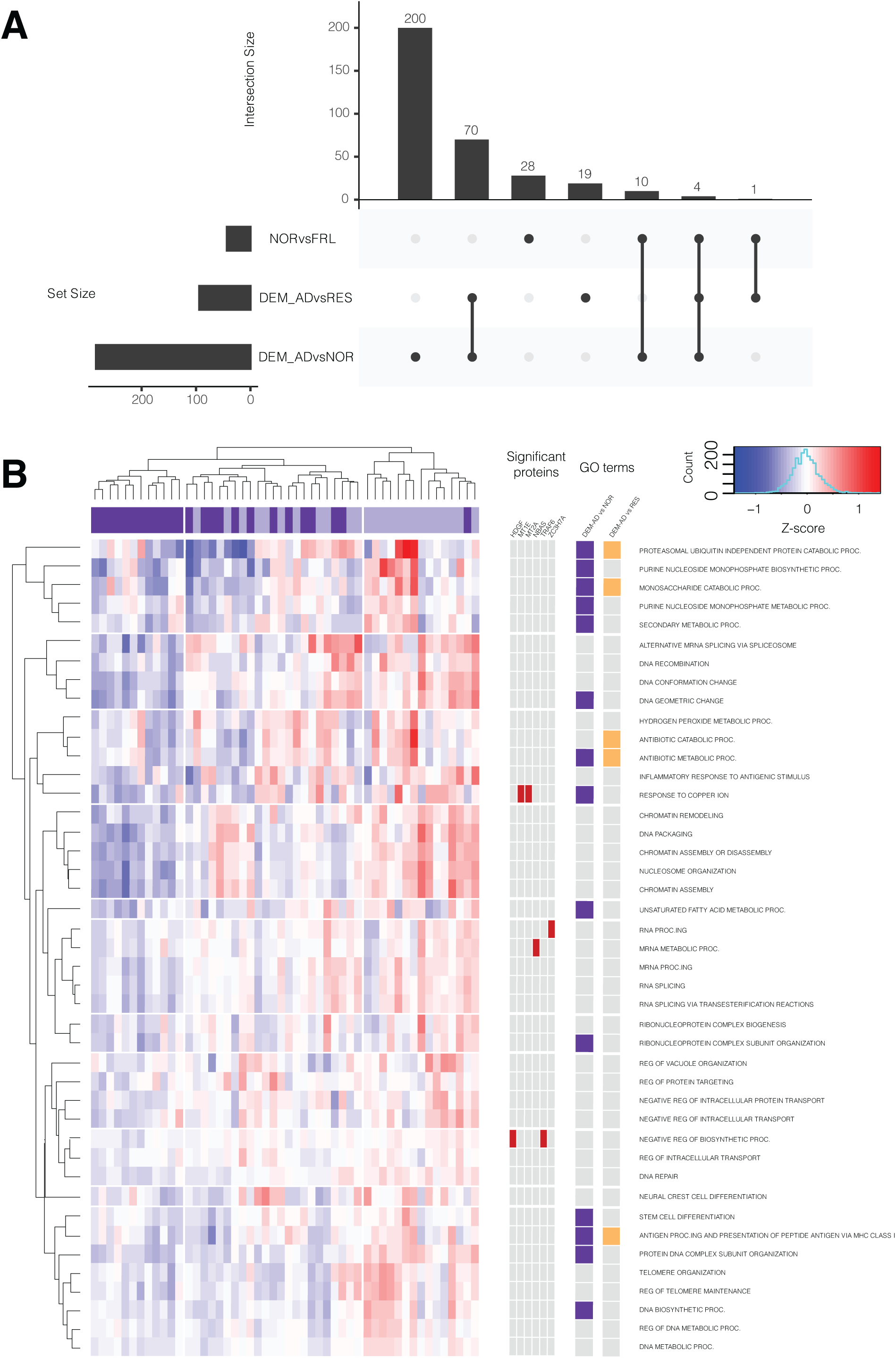
Summary of Gene Set Enrichment Analysis from cognitively contrasted samples shows only a small overlap between terms related to Frailty and terms related to Resilience. **A)** Upset plot shows the number of GO terms shared between each cognitive contrast. The majority of the terms that are significant in the Normal vs Frail contrast are unique to this contrast. **B)** Heatmap shows FDR significant (p < 0.05) clustered GO-terms from the Frail versus Normal comparison. Significant proteins in this contrast are shown alongside the GO terms they are associated with. Comparison with the significant terms from other cognitive contrasts is shown alongside. Common words in GO terms have been shortened for plotting; Proc. = Process, Reg. = Regulation, Sig. = Signaling.

The Dementia-AD versus Resilient comparison more closely reflected that of the Dementia-AD versus Normal comparison (Table S11, Figure 6A). 94 GO terms were significant in this contrast, of which 74 were shared with the Dementia-AD versus Normal contrast. Nineteen terms were unique to this contrast, including a small cluster of three terms related to dopamine metabolism, and a cluster of four Humoral Immune response terms. Both of these terms were enriched in Dementia-AD samples compared to Resilient. Terms related to Glutamate signaling and memory formation were enriched in Resilient samples compared to Dementia-AD, despite there being no gross reduction in established synapse markers in Dementia-AD. Terms for mitochondrial oxidative phosphorylation were also enriched in Resilient. In the opposite direction, metabolic processes such as carbohydrate metabolism were enriched in Dementia-AD compared to Resilient samples. Immune categories, mostly related to antigen presentation, were also enriched in the Dementia-AD samples compared to the Resilient samples. Clustering of these GO terms did not strongly separate Dementia-AD from Resilient samples, suggesting that it is more difficult to define these two groups at the level of synaptic proteins (Figure 7). At the protein level CHGA and SEMA7A, both more abundant in Dementia-AD, are associated with Immune Activation related GO terms, while SLC8A3, upregulated in Resilience, is associated with Regulation of Synaptic Plasticity.

**Figure 7:**
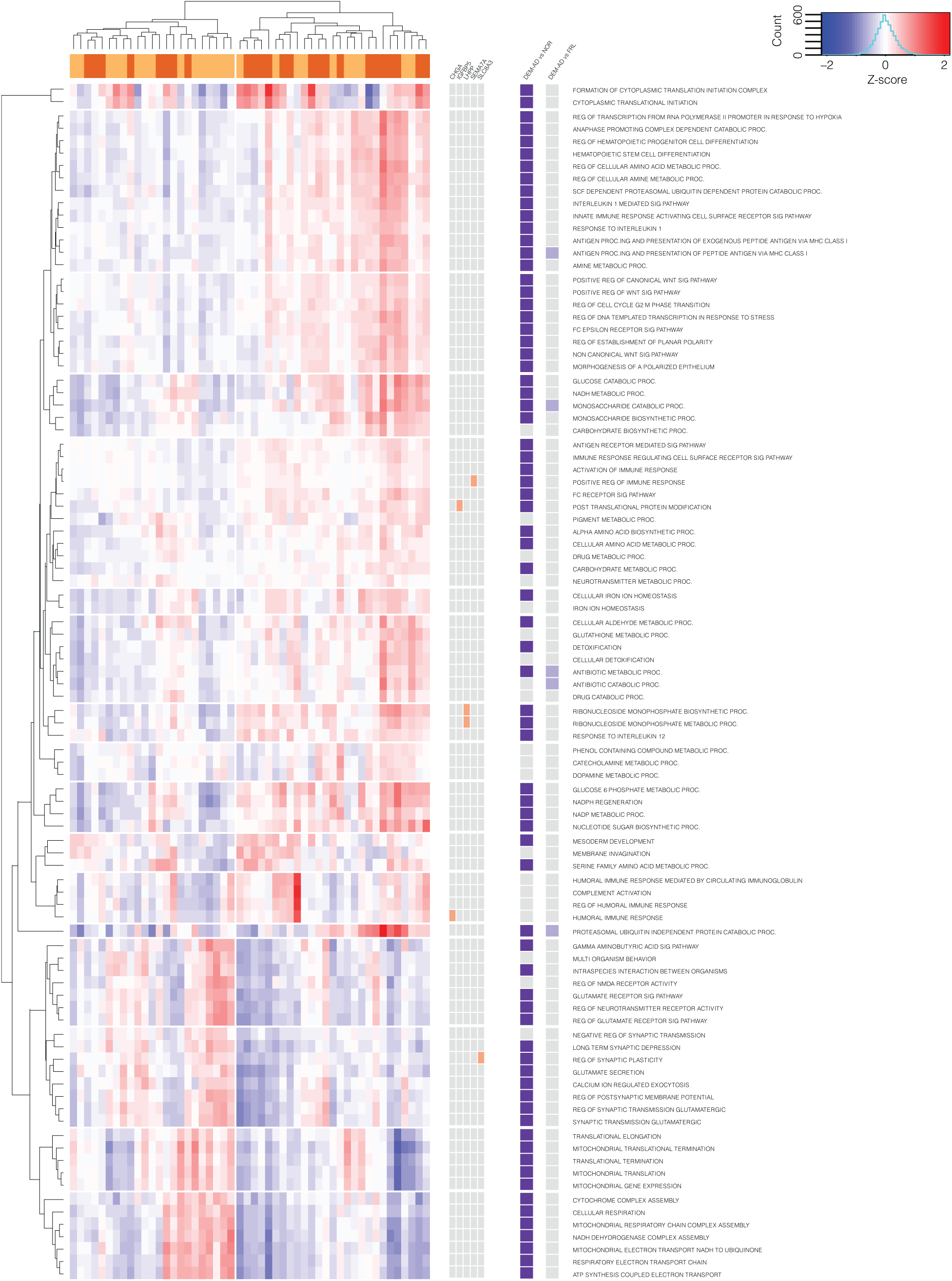
Summary of Gene Set Enrichment Analysis from Dementia-AD versus Resilient samples shows substantial overlap with the Dementia-AD versus Normal contrast. Heatmap shows FDR significant (p < 0.05) clustered GO-terms from the Dementia-AD versus Resilient comparison. Significant proteins in this contrast are shown alongside the GO terms they are associated with. There is substantial overlap between these GO terms and those significant in the Dementia-AD versus Normal comparison. Common words in GO terms have been shortened for plotting; Proc. = Process, Reg. = Regulation, Sig. = Signaling

**Figure 8:**
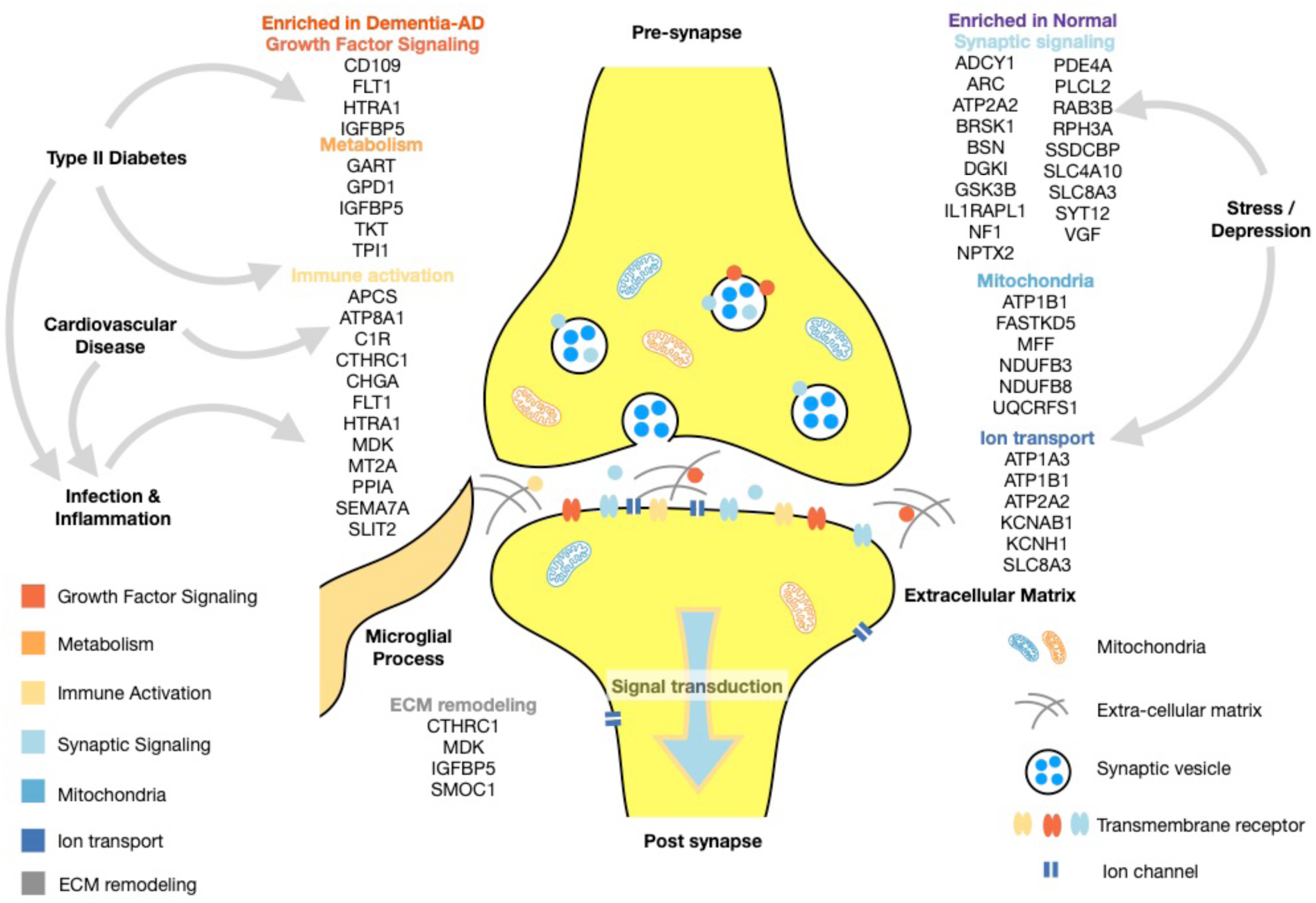
Schematic overview highlighting key proteins and functional categories.

## Discussion

Although the presence of substantial amyloid-β plaques is required for a diagnosis of AD, studies of community residing older adults have shown that up to one third of older people with no cognitive impairment at death harbor neuropathology that would be classified as intermediate to high likelihood of AD (D. A. Bennett et al., 2006; Schneider et al., 2007). Amyloid-β plaque presence is therefore necessary but not sufficient for causing the cognitive manifestations of AD. General synapse loss has long been suggested as a strong predictor of cognitive decline in AD (DeKosky & Scheff, 1990; Koffie et al., 2011; Terry et al., 1991), as reflected in whole tissue studies of AD (Johnson et al., 2018b; Ping et al., 2018; Wingo et al., 2019). To avoid this volume effect-confound, mass spectrometry (LC-MS3) was used to provide a more comprehensive, detailed and unbiased examination of the proteomes of enriched synapses. This enabled identification of protein changes within synapses that were associated with cognitive performance. To model the association of synapse proteins with cognitive performance and AD pathology as independent variables, we chose an experimental structure with four distinct diagnostic groups which were well matched for key sample demographics. Alongside the highest contrast comparison of Dementia-AD and Normal cases, we also included two clinicopathologically discordant groups: Resilient subjects with high AD pathology burden but cognitive resilience, and Frail subjects with low AD pathology burden but impaired cognitive performance. AD is a complex disease driven by multiple pathophysiologies, including protein misfolding (Chiti & Dobson, 2017), inflammation (Heppner, Ransohoff, & Becher, 2015; Salter & Stevens, 2017), oxidative injury (Kim, Kim, Rhie, & Yoon, 2015), metabolic disturbances (Arnold et al., 2018; Ribe & Lovestone, 2016), and neurovascular dysfunction. Progressive dementia is the clinical manifestation of these processes that insidiously evolve over years prior to the expression of clinical symptoms. While whole tissue proteomic studies regularly highlight inflammatory and mitochondrial dysregulation as key proteomic changes in AD tissue, with the former suggested to reflect the activation state of glial cells in whole tissue, it is currently unclear exactly how these processes interact to affect the synapse, the cellular compartment most directly related to cognitive dysfunction in AD.

Gene Set Enrichment Analysis of our study showed that GO terms related to immune function, NAD and Glucose metabolism, and ECM remodeling were all more prevalent in synaptic proteins from the Dementia-AD samples compared to Normal samples (Figure 7). This establishes the synapse as a potential site of metabolic dysfunction in cases of impaired cognition, regardless of the presence of significant AD pathology. Changes to the NADH/NAD+ balance in aging tissue have been implicated in AD pathophysiology, and may differ between the mitochondria and the cytoplasm (Hou et al., 2018; Stein & Imai, 2012). This balance is the target of a novel therapeutic for AD, nicotinamide riboside, which has shown preclinical efficacy in multiple mouse models of AD (Gong et al., 2013; Hou et al., 2018) and a single trial of human Parkinson’s Disease (Birkmayer, Vrecko, Volc, & Birkmayer, 1993). GK, PDHA1, PDK3, PPIP5K2, and are all enzymes involved in carbohydrate metabolism which are decreased in Dementia-AD synaptic samples in our study. Regional glucose utilization is decreased in AD subjects as evidenced by decreased signals seen with FDG PET, and this especially so in the angular gyrus (Vlassenko & Raichle, 2015). This dysregulation may add weight to the notion that the synapse is a critical site for the effects of Type II Diabetes as a risk factor for dementia, brain insulin resistance and metabolic dysregulation (Arnold et al., 2018).

Immune and inflammation related terms were also enriched in all Dementia-AD contrasts, in agreement with previous whole tissue studies (Johnson et al., 2018a; Wingo et al., 2019; Zhang et al., 2018). Humoral response GO terms were unique to the Dementia-AD versus Resilience comparison, being elevated in Dementia-AD. Protein drivers of this category APCS, CHGA, C1R, IGHA1, and IGLL5 are all enriched in Dementia-AD. Complement signaling (C1R) has been shown to be active at the synapse, with the later cascade component C4A known to localize to the pre- and post-synapse, dendrites, and axons. C4A has been genetically and mechanistically implicated in synaptic dysfunction in both schizophrenia (Sekar et al., 2016) and AD (Hong et al., 2016; Zorzetto et al., 2016). The complement system is often shown to be dysregulated in proteomic studies of plasma in AD (B. Carlyle, Trombetta, & Arnold, 2018)56), suggesting this system as a potential point of cross talk between peripheral inflammatory risk factors and AD (Kinney et al., 2018). While synaptic localization for C4A is clear from multiple studies, other proteins driving enrichment of the immune categories are less well defined, and may be arising from the incursion of glia, particularly microglia, into the fractionated synaptic cleft.

In agreement with previous studies (Johnson et al., 2018a; Wingo et al., 2019), cognitively Resilient and Normal samples showed strong enrichment for both synaptic signaling and mitochondrial GO terms. However, the hub proteins driving these categories in whole tissue studies are more strongly reflective of gross synapse loss in AD samples, including PSD95, SYT1 and STX1A. In our study, we designed our approach to minimize the effect of bulk loss of synapses, and did not see any between group differences in these synaptic marker proteins. We were therefore able to see changes within synapses beyond this gross synaptic loss. We highlighted a different group of proteins that were maintained in cognitively normal samples compared to cognitively impaired, including the immediate early protein ARC, RPH3A which is involved in calcium dependent exocytic release (Tan et al., 2014) and GluN2A PSD95 interactions (Stanic et al., 2015), the membrane trafficking SYT12, the neuropeptide VGF and the adenylyl cyclase pathway molecules ADCY1 and PDE4A, and KCN channel components. Adenylyl cyclase signaling and regulation of membrane potential downstream of PDE4A, is involved in regulation of cAMP signaling in higher order cortical circuits in primates (B.C. Carlyle et al., 2014). This carefully balanced signaling pathway is exquisitely sensitive to stress, and may be interrupted in AD (Becky C Carlyle et al., 2014). Mitochondrial categories were enriched for mitochondrial respiratory chain components (NDUF proteins). This is likely a reflection of the number of appropriately sized mitochondria present in the pre- and post-synapse. Mitochondrial numbers and morphology are regulated by a complicated balance of fusion and fission known to be disrupted in AD (Flannery & Trushina, 2019). Electron microscopy has shown decreased pre- and post-synaptic mitochondria in the superior temporal gyrus in human post-mortem tissue (Pickett et al., 2018).

While there are a number of proteins that clearly track with the presence of AD neuropathology in every pathological contrast (CTHRC1, SLIT2, CPSF6, FLT1, SMOC1, MDK, and SNRNP70), there are no proteins and only four GO terms (Proteasomal Ubiquitin Independent Protein Catabolic Process, Monosaccharide Catabolic Process, Antibiotic Metabolic Process and Antigen Processing and Presentation of Peptide Antigen) that unite all cognitive contrasts. This may simply be a result of variability between individuals – amyloid-β plaque and tau tangle environments may have a relatively consistent protein signature between individuals, whereas there are multiple different pathways that may result in neurodegenerative dementia. Subtle dysregulations in metabolic, immune, or stress pathways over a lifetime may interact at the synapse to produce signaling dysfunction that are the initial changes associated with memory impairment. This is especially true for the Frail group, where none of the significantly upregulated proteins have been well studied in the CNS. The Frail group was not enriched for any other gross or microscopic neuropathological features (Figure S2), so this is likely not a reflection of mixed pathology dementia. It further appears that cognitive resilience in the face of AD pathology is not simply the opposite of frailty in the absence of gross pathology, but that resilient subjects maintain key synaptic signaling regulator proteins and mitochondrial proteins despite the presence of plaques, tangles and the immune upregulation that associates with them.

Though proteomics is still a relatively nascent field, it is clear that studies of post-mortem human AD brain tissue, in conjunction with a generally open attitude to sharing of data and subject metadata, are leading to converging findings on a number of proteins that were not previously thought to be associated with AD pathophysiology. VGF, IGFBP5, NPTX2, SMOC1 and MDK are clear examples of converging data, appearing in multiple studies despite having received very little attention in the field prior to the widespread use of proteomics. Sample enrichment, such as performed here, can uncover new targets such as LHPP, CPSF6, MACROD1, SEMA7A, and SNRNP70 due to an increased sensitivity to changes in more localized regions, cellular and subcellular compartments. Through this approach we have highlighted novel proteins at the synapse that may be involved in the intersecting pathophysiologies that drive AD progression (Figure 7), and which may represent novel therapeutic targets or biomarkers of engagement for medications targeting inflammation, mitochondrial function, and synaptic modulation.

## Materials and Methods

### Human brain tissue

Post-mortem tissue from the parietal association cortex (angular gyrus, Brodmann Area 39) was obtained from the Rush Alzheimer’s Disease Center. Tissue came from both the Religious Orders Study (ROS) and the Memory and Aging Project (MAP) (David A Bennett et al., 2018), similarly designed studies with longitudinal cohorts consisting of individuals who agreed to annual clinical evaluations and provided informed consent to donate their brains for research at the time of death. Annual evaluations included a medical history, neurological exam, and twenty-one cognitive tests assessing multiple cognitive domains that are commonly impaired in older individuals (Wilson, Bienias, Evans, & Bennett, 2004; Wilson et al., 2002). Cognitive scores were converted to Z-scores across the entire ROSMAP cohort and combined to generate a composite global cognition score. Brain autopsies were conducted with standardized protocols, including the preparation of diagnostic blocks for neuropathological classification according to NIA-Reagan, Braak, and CERAD staging. Case metadata is provided in Supplementary Table 1. Tissue was obtained and analyzed under an Exempt Secondary Use protocol approved by the Massachusetts General Hospital Institutional Review Board (2016P001074).

### Case selection and categorical grouping

In total, 100 cases that spanned the range of AD pathology and last valid global cognition scores in the ROSMAP cohorts were selected. These 100 cases were selected from 4 diagnostic bins stratified based on two variables. The first was the Braak Score, a 6 point scale that describes the brain-region specific pattern of AD pathology from early to late stage disease. Subjects were divided into low AD pathology (Braak Score of 4 or less) or high AD pathology (Braak Score of 5 or 6) on the basis of this variable. The second variable was a consensus clinical impression from longitudinal cohort clinicians as to whether the subject was cognitively impaired at death. By dividing cases on the basis of these two variables, four groups were selected with n = 25 per group. Selected cases in categorical groups were well balanced for other key demographic variables including age, gender, education and post-mortem interval (Table 1).

### Synaptic protein enrichment

Syn-PER Synaptic Protein Extraction Reagent (Thermo Fisher Scientific, Waltham, MA, USA) was used to enrich for synaptic proteins from the frozen tissue samples. Complete Protease Inhibitor (EDTA Free, Roche, Mannheim, Germany) was added to the Syn-PER reagent (1 tablet per 50 ml). After removal of obvious white matter, approximately 100 mg of each frozen tissue piece was weighed, before adding 1 ml of Syn-PER per 100 mg of tissue and homogenization using 15 strokes of a dounce homogenizer driven by a Scilogex (Rocky Hill, CT, USA) OS-20S drive motor set at 500 rpm. Homogenates were centrifuged at 1,200 x g for 10 min at 4°C. The supernatant (S1) was transferred to a new sample tube and centrifuged at 15,000 x g for 20 min at 4°C. The supernatant (S2) was removed and discarded, and the resulting P2 pellet was resuspended once in cold PBS to reduce contaminating proteins, before a second spin at 15,000 x g for 20 min at 4°C. The washed P2 pellet was snap frozen and stored at −80 °C until it was processed for analysis by TMT-LC-MS3.

### Multiplexed quantitative proteomics

P2 pellets were lysed by passing through a 21-gauge needle twenty times in 75 mM NaCl, 3% SDS, 1 mM NaF, 1 mM beta-glycerophosphate, 1 mM sodium orthovanadate, 10 mM sodium pyrophosphate, 1 mM PMSF and 1x Roche Complete Mini EDTA free protease inhibitors in 50 mM HEPES, pH 8.5. Lysates were then sonicated for 5 min in a sonicating water bath before cellular debris was pelleted by centrifugation at 14000 rpm for 5 min. Proteins were then reduced with DTT and alkylated with iodoacetamide as previously described (Edwards & Haas, 2016) and purified through methanol-chloroform precipitation (Wessel & Flügge, 1984). Precipitated proteins were reconstituted in 1 M urea in 50 mM HEPES, pH 8.5, digested with Lys-C and trypsin, and desalted using C18 solid-phase extraction (SPE) (Sep-Pak, Waters, Beverly, MA, USA). The concentration of the desalted peptide solutions was measured by BCA assay, and peptides were aliquoted into 50 μg portions. Peptide samples were randomized and labeled with TMT11 as described previously (Edwards & Haas, 2016). They were pooled into sets of ten samples and a bridge sample generated from mixing parts of the digests of all sample was added to each pool (Lapek et al., 2017). The pooled samples were desalted via C18 SPE and fractionated using Basic pH Reversed-Phase Liquid Chromatography (bRPLC) (Edwards & Haas, 2016).

Twelve fractions were analyzed by LC-MS2-MS3 on an Orbitrap Fusion mass spectrometer (Thermo Fisher Scientific, Waltham, MA, USA) coupled to an Easy-nLC 1000 autosampler and HPLC system. Peptides were separated on an in-house pulled, in-house packed microcapillary column (inner diameter, 100 μm; outer diameter, 360 μm, 30 cm GP-C18, 1.8 μm, 120 Å, Sepax Technologies, Newark, DE, USA). Peptides were eluted with a linear gradient from 11 to 30% ACN in 0.125% formic acid over 165 minutes at a flow rate of 300 nL/minute while the column was heated to 60 °C. Electrospray ionization was achieved by applying 1500 V through a stainless-steel T-junction at the inlet of the microcapillary column.

The Orbitrap Fusion was operated in data-dependent mode using an LC-MS2/SPS-MS3 method. Full MS spectra were generated over an m/z range of 500-1200 at a resolution of 6 × 10^4^ with an AGC setting of 5 × 10^5^ and a maximum ion accumulation time of 100 ms. The most abundant ions detected in the survey scan were subjected to MS2 and MS3 experiments using the Top Speed setting that enables a maximum number of spectra to be acquired in a 5 second experimental cycle before the next cycle is initiated with another survey full-MS scan. Ions for MS2 spectra were isolated in the quadrupole (0.5 m/z window), Collision Induced Dissociation (CID)-fragmented, and analyzed at rapid scan rate in the ion trap, where fragment ions were analyzed (Automatic Gain Control (AGC), 10 × 10^4^; maximum ion accumulation time, 35 ms; normalized collision energy, 30%). MS3 analysis was performed using synchronous precursor selection (SPS MS3) upon Higher Energy Collisional (HCD) fragmentation. Up to 10 MS2 precursors were simultaneously isolated and fragmented for MS3 analysis (isolation width, 2.5 m/z; AGC, 1 × 10^5^; maximum ion accumulation time, 100 ms; normalized collision energy, 55%; resolution, 6 × 10^4^). Fragment ions in the MS2 spectra with an m/z of 40 m/z below and 15 m/z above the precursor m/z were excluded from being selected for MS3 analysis.

### Data processing

Data were processed using an in-house developed software suite (Huttlin et al., 2010b). RAW files were converted into the mzXML format using a modified version of ReAdW.exe (http://www.ionsource.com/functional_reviews/readw/t2x_update_readw.htm). Spectral assignments of MS2 data were made using the Sequest algorithm (Eng et al., 1994) to search the Uniprot database (02/04/2014 release) of human protein sequences including known contaminants such as trypsin. The database included a decoy database consisting of all protein sequences in reverse order (Elias & Gygi, 2007). Searches were performed with a 50 ppm precursor mass tolerance. Static modifications included 11-plex TMT tags on lysine residues and peptide n-termini (+229.162932 Da) and carbamidomethylation of cysteines (+57.02146 Da). Oxidation of methionine (+15.99492 Da) was included as a variable modification. Data were filtered to a peptide and protein false discovery rate of less than 1% using the target-decoy search strategy (Elias & Gygi, 2007). This was achieved by first applying a linear discriminator analysis to filter peptide annotations using a combined score from the following peptide and spectral properties (Huttlin et al., 2010a): XCorr, ΔCn, missed tryptic cleavages, peptide mass accuracy, and peptide length. The probability of a peptide-spectral match to be correct was calculated using a posterior error histogram and the probabilities of all peptides assigned to one specific protein were combined through multiplication. The dataset was re-filtered to a protein assignment FDR of less than 1% for the entire dataset of all proteins identified across all analyzed samples (Huttlin et al., 2010b). Peptides that matched to more than one protein were assigned to that protein containing the largest number of matched redundant peptide sequences following the law of parsimony (Huttlin et al., 2010b).

For quantitative analysis, TMT reporter ion intensities were extracted from the MS3 spectra by selecting the most intense ion within a 0.003 m/z window centered at the predicted m/z value for each reporter ion, and signal-to-noise (S/N) values were extracted from the RAW files. Spectra were used for quantification if the sum of the S/N values of all reporter ions was ≥ 440 and the isolation specificity for the precursor ion was ≥ 0.75. Protein intensities were calculated by summing the TMT reporter ions for all peptides assigned to a protein. Normalization of the quantitative data followed a multi-step process. Intensities were first normalized using the intensity measured for the bridge sample (Lapek et al., 2017). Taking account of slightly different protein amounts analyzed in each TMT channel, we then added an additional normalization step by normalizing the protein intensities measured for each sample by the median of the median protein intensities measured in these samples.

### Data analysis

Median normalized protein quantifications were imported into R for all downstream analysis. All code used from this point forwards is provided in R project format, including non-PDF format supplementary tables. In building protein fraction consensus lists, protein ID conversion was performed using downloaded flat files from biomart (Smedley et al., 2015) (Mouse version: GRCm38.p6, Human version: GRCH38.p12) when necessary. Due to wide variation in outlying values, data was further normalized by quantile normalization in the PreProcessCore R package. Linear modelling, ANOVAs, Tukey HSD tests, and correction for multiple comparisons were performed using base R functions. The ggsignif and scales packages were used in combination with ggplot for data plots. Files for input to GSEA (Subramanian et al., 2005) were prepared using a base R script. GSEA analyses were run using the java applet downloaded from http://software.broadinstitute.org/gsea/index.jsp, with downstream analysis and plotting performed in R. Ontological trees were collapsed using ontological information from the GSA package.

### Data availability

For initial review purposes, supplementary tables may be accessed through this dropbox link: https://www.dropbox.com/sh/gdsrf1y9yr6xbia/AAA6XwvTio912q_Mcv-xBo1fa?dl=0

“Mass spectrometry RAW data are accessible through the MassIVE data repository (massive.ucsd.edu) under the accession number MSV000084959.

These data will be made public as soon as the paper is accepted.

For reviewer access please provide the following username and password:

Username: MSV000084959_reviewer

Password: Carlyle

The code used to create all analyses, figures and supplementary tables for this manuscript can be found at: https://bitbucket.org/omicskitchen/tmtsynaptosomes/src/master/

Individual protein abundances across key variables can be explored at https://tmt-synaptosomes.omics.kitchen/

## Acknowledgements

This work was supported by the Challenger Foundation / Minehan-Corrigan Family and by NIH R01 AG039478, R01 AG062306, P30 062421, R01 AG017917, P50 AG047270, R01 AG15819, P30 AG20262, R01 AG15819, R01 AG17917, and U01 AG61356. BCC is supported by the Bright Focus Foundation.

---

**Figure S1:**
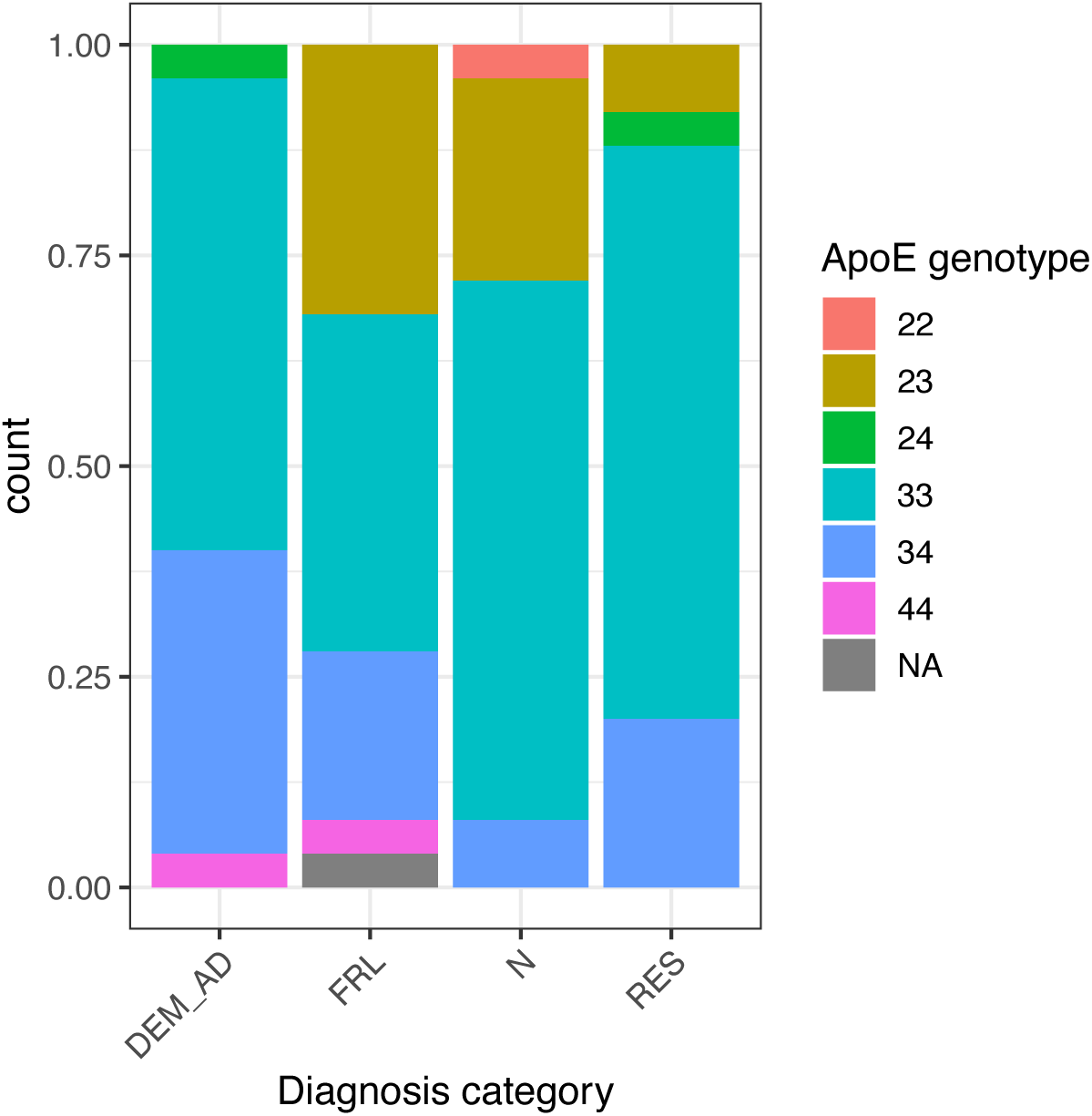
ApoE allele distribution was not matched between the four groups. Stacked bar plot showing ApoE genotype distribution by group. There were the most ApoE4 risk allele carriers in the Dementia-AD group. Interestingly, the Frail group had the largest number of ApoE2 protective allele carriers. Data was unavailable for one Frail subject.

**Figure S2:**
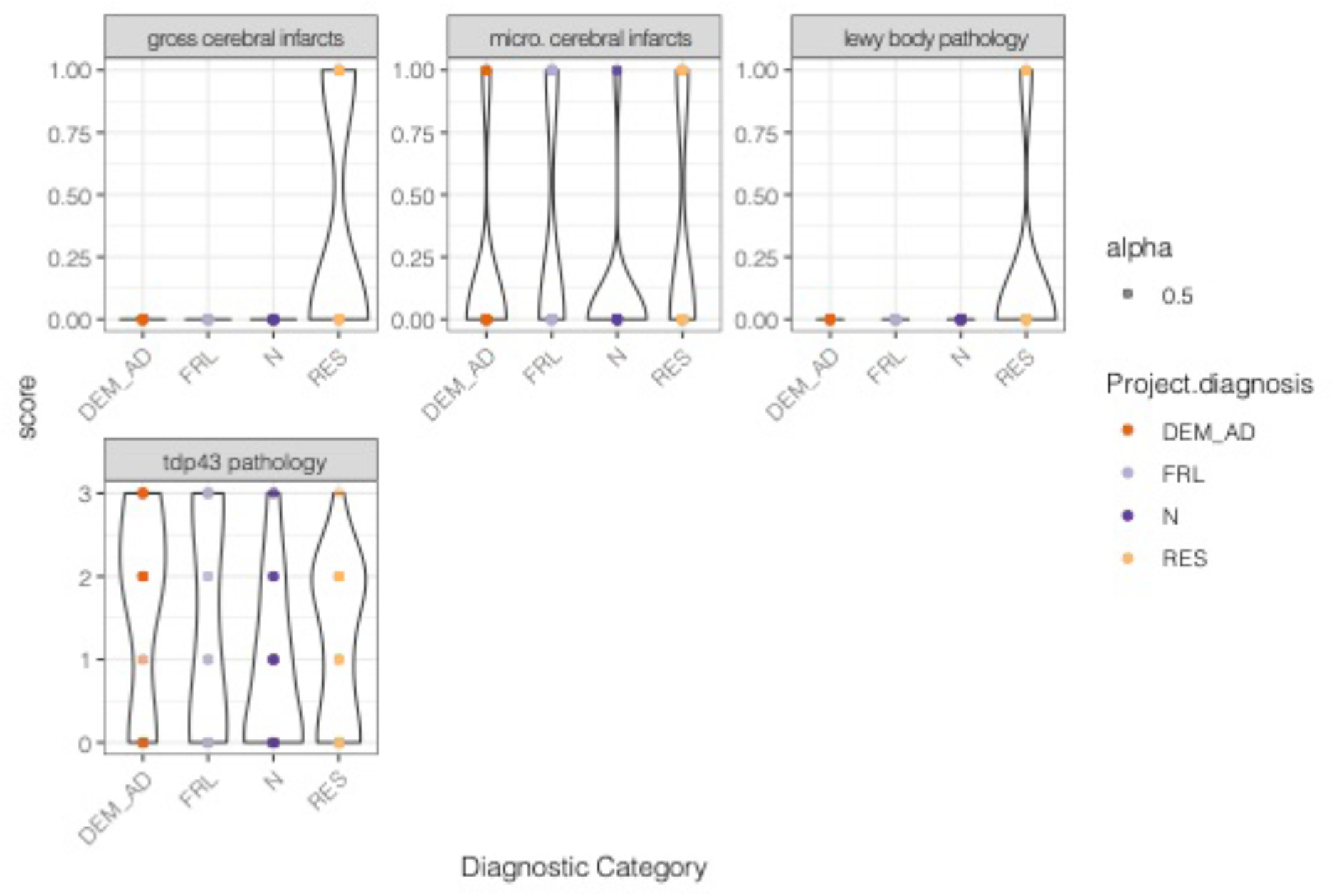
Other gross and microscopic pathologies were generally evenly spread across groups. Gross & microscopic vascular infarcts and lewy body pathology, detailed as present (1) or absent (0). TDP43 pathology is ranked in four stages, with 0 being no TDP43 pathology and 3 being amygdala, limbic and cortical involvement. There were 16 missing values for TDP43 pathology.

**Figure S3:**
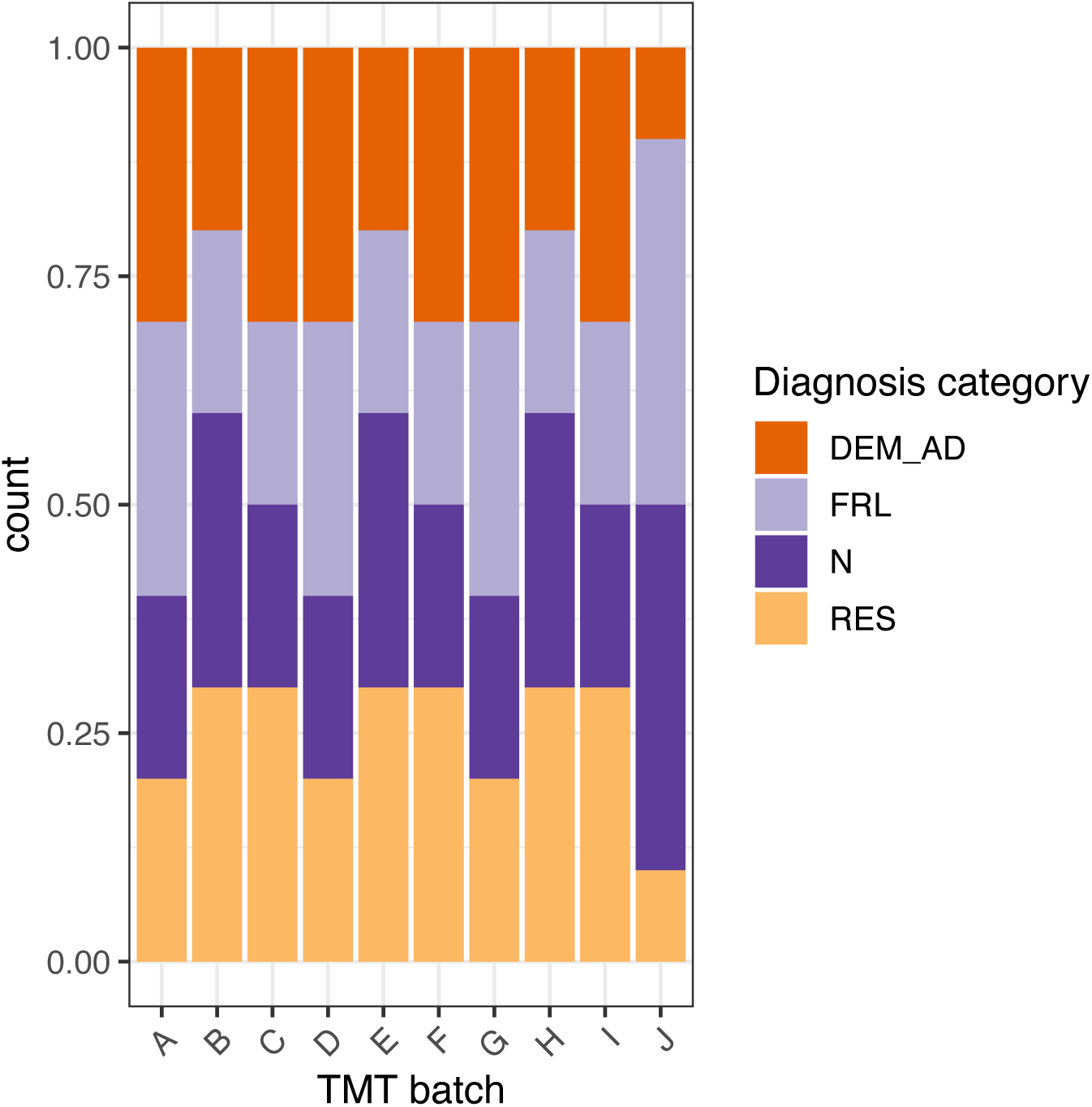
Diagnostic conditions are well balanced across the 10 TMT 10-plex batches. Bar plot shows the number of samples from each diagnostic condition across all ten batches A to J.

**Figure S4:**
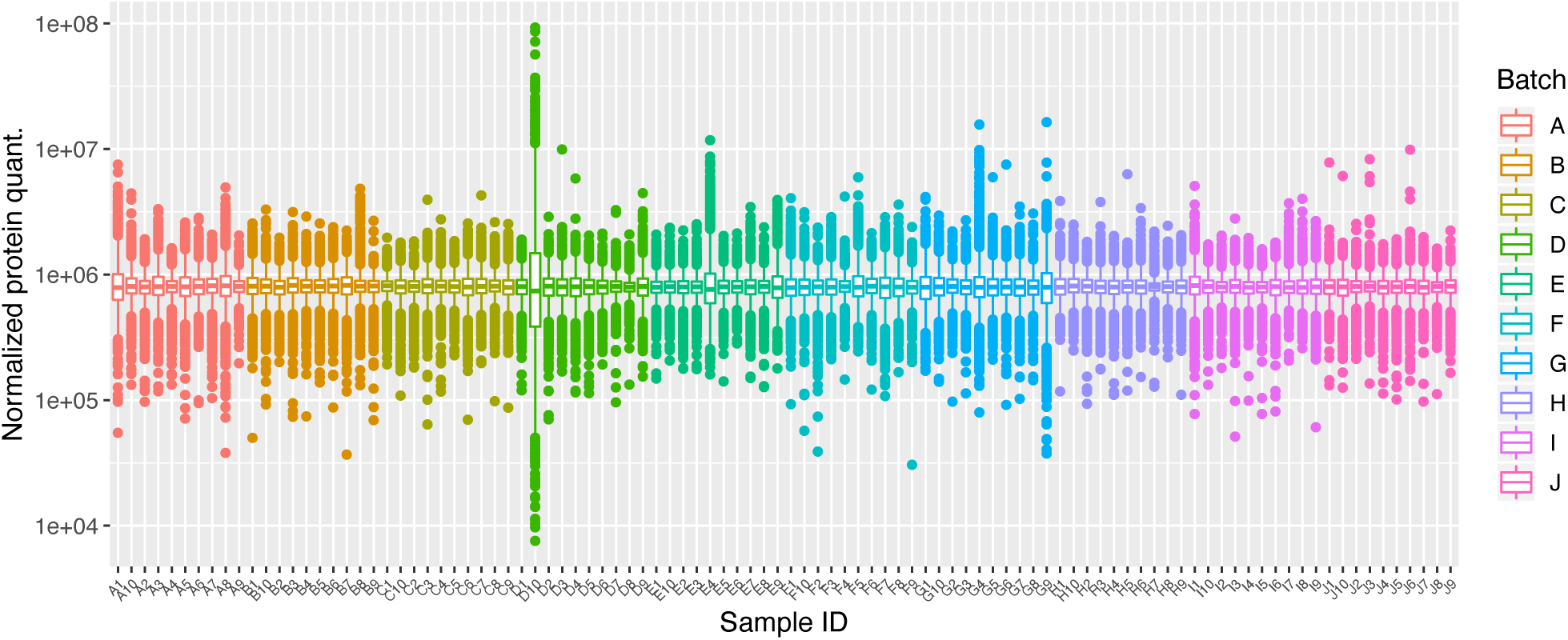
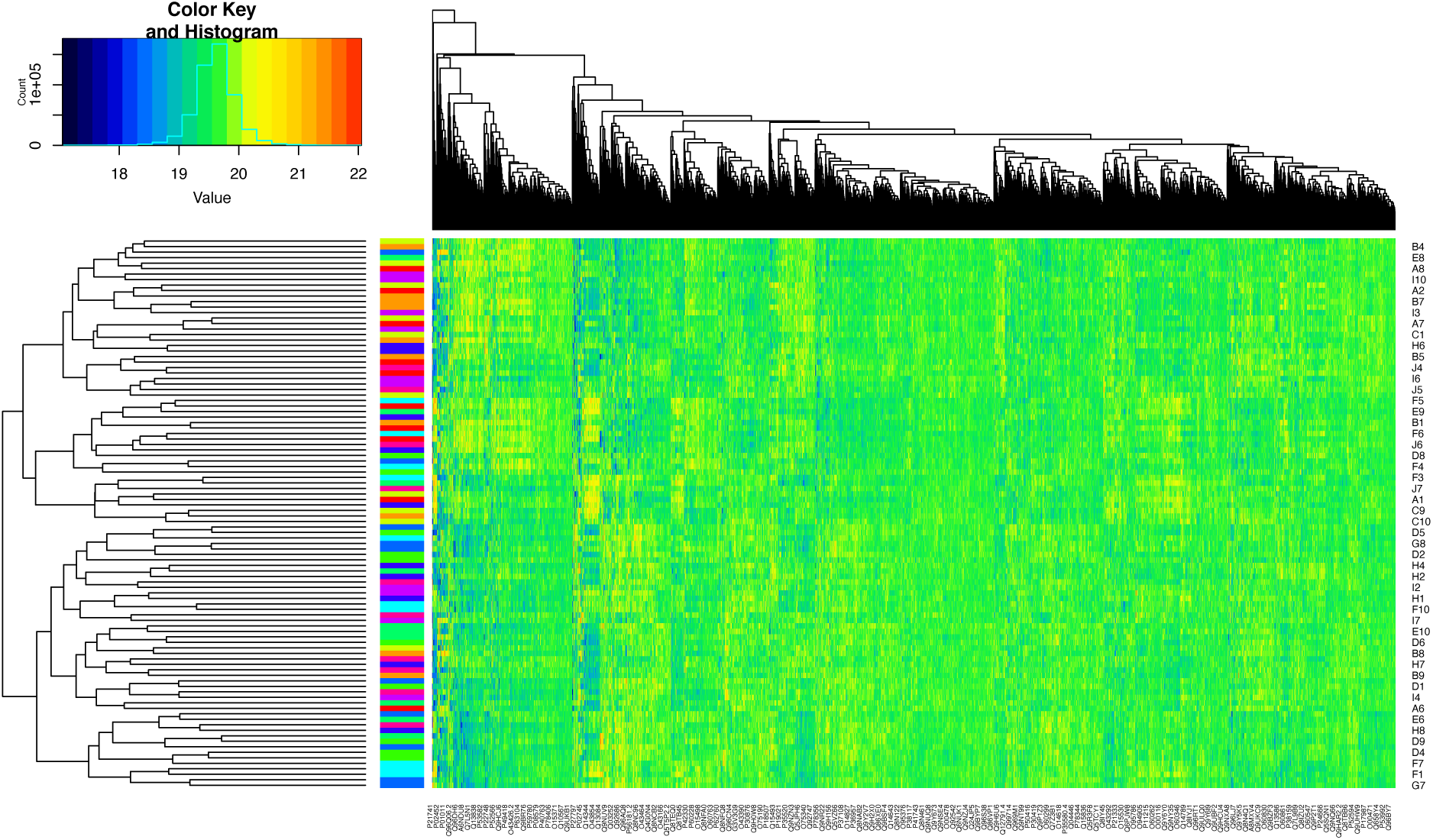
Quality control plots show effective normalization of data. **A)** Box plots show appropriate two-step median normalization across TMT 10-plex batches. In these blots the box indicates the bounds of the 25th and 75th percentile, with the central line representing the median. The whiskers extend to 1.5 times the inter quartile range, and individual proteins outside these limits are plotted as single point outliers. **B)** Heatmap and hierarchical clustering shows no obvious signs of batch effects arising from TMT 10-plexes after quartile normalization. Row side labels are color coded by batch

**Figure S5:**
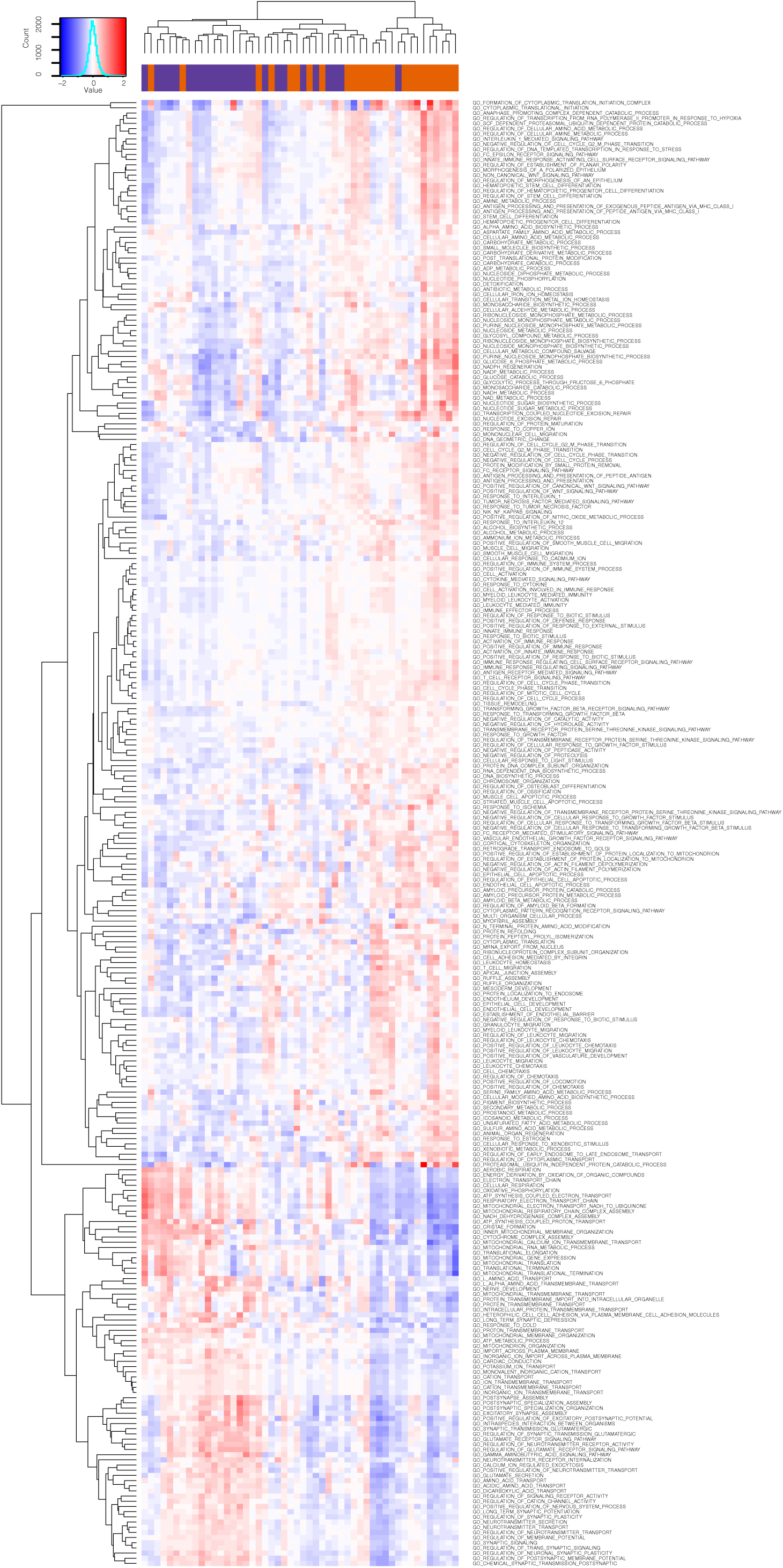
284 GO terms are significant by Gene Set Enrichment Analysis (GSEA) in the Normal versus Dementia-AD comparison. This zoomable figure shows all 284 terms without any collapsing by parent term.

**Figure S6:**
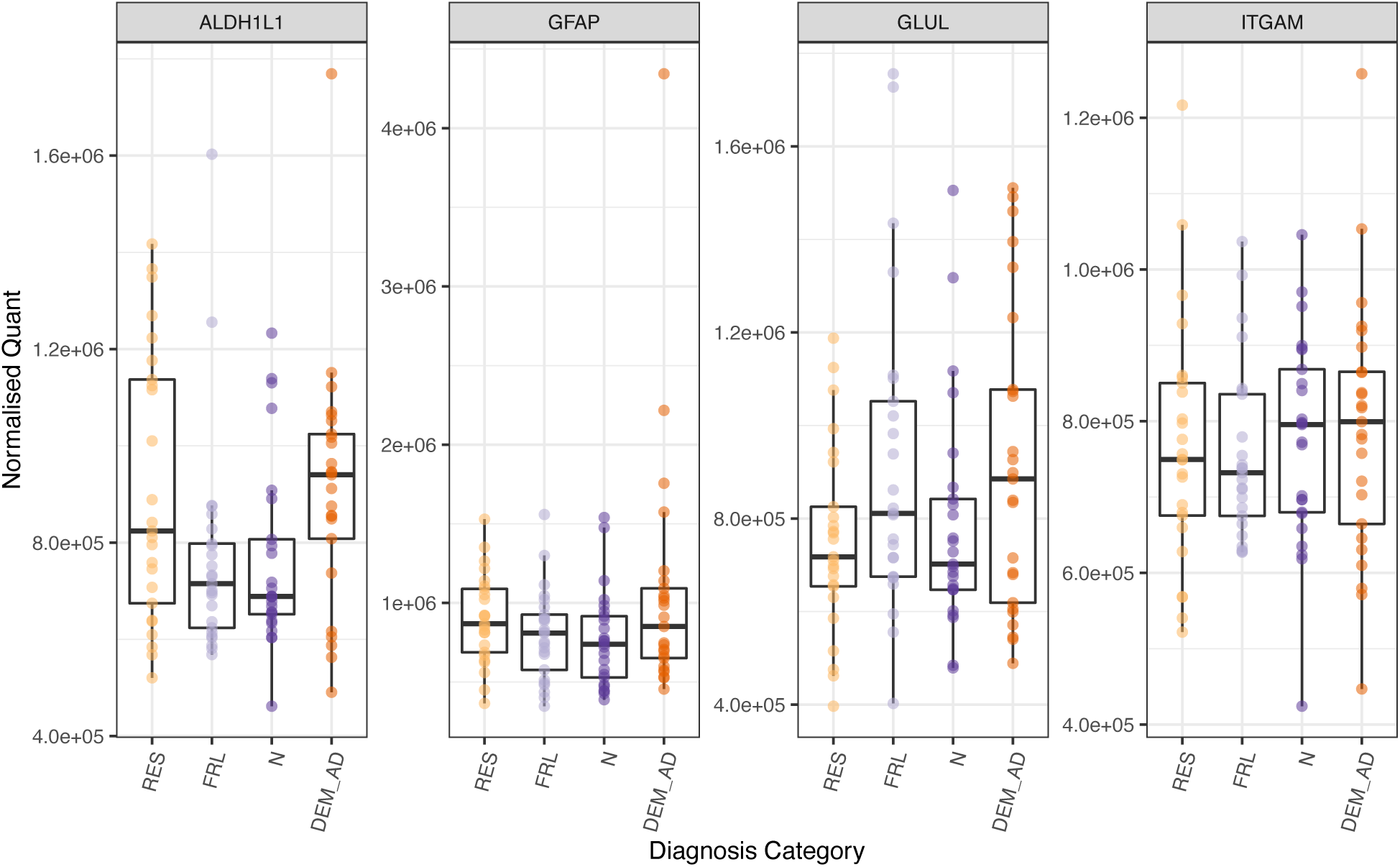
There are no significant between group differences in Astrocyte (ALDH1L1, GFAP, GLUL) or Microglial markers (ITGAM).

## Supplementary Data

Table S1: Individual sample metadata

Table S2: Summary of Tukey test outcomes from demographic data

Table S3: Table summarizing sub cellular fraction enrichment Fisher exact tests.

Table S4: Synaptic index ANOVA data

Table S5: Table showing status of all proteins across all terms in the linear modeling

Table S6: Summary of significant proteins from the linear modelling, all variables included in one table

Table S7: GSEA GO category enrichment results for the Normal vs Dementia-AD comparison

Table S8: GSEA GO category enrichment of significant proteins in the Normal vs Dementia-AD comparison

Table S9: GSEA GO category enrichment results for the Normal vs Frail Comparison

Table S10: GSEA GO category enrichment of significant proteins in the Normal vs Frail comparison

Table S11: GSEA GO category enrichment results for the Resilient vs Dementia-AD Comparison

Table S12: GSEA GO category enrichment of significant proteins in the Resilient vs Dementia-AD Comparison

